# Interpreting Graph Neural Networks for Antibacterial Drug Repurposing against *Acinetobacter baumannii*

**DOI:** 10.64898/2026.10.26.761825

**Authors:** Salvatore Romano, Marco Grassia, Dario Righelli, Pietro Liò, Giuseppe Mangioni, Cristian Taccioli

## Abstract

Antimicrobial resistance in *Acinetobacter baumannii*, whose carbapenem-resistant strains are classified by the World Health Organization as a critical priority, has motivated machine-learning screens for new antibacterials, but translating predictions into chemical hypotheses requires evaluating the molecular features on which a model relies. A graph neural network ensemble achieves competitive predictive performance against a descriptor-augmented reference model on the same antibacterial screen while using only atom features and connectivity, allowing interpretation to focus on the molecular graph without a separate input of handcrafted descriptors. The atom-level explanations identify features supporting the model’s activity predictions under perturbation and restoration, while comparisons with equally sized perturbations within each molecule distinguish piperazine as the most consistently supported recurring contributor to predicted activity. Prospective screening of the Drug Repurposing Hub followed by developability and safety assessment yields a ranked shortlist for experimental follow-up and leads to testing of CE3F4, which reduced *A. baumannii* growth by 57% at 0.25 mM in a preliminary assay with a non-monotonic concentration–response profile.

## 1 Introduction

Antimicrobial resistance is associated with substantial global mortality [1] and can leave few effective treatment options for infections caused by multidrug-resistant strains [2, 3]. Among the pathogens contributing to this burden, *Acinetobacter baumannii* is a Gram-negative bacterium that persists in health-care environments and causes severe hospital-acquired infections [4]. These treatment limitations and the limited innovation in the antibacterial development pipeline [5] motivate complementary discovery strategies such as drug repurposing [6].

Phenotypic screening can identify repurposing candidates by testing existing compounds for antibacterial activity without requiring a predefined molecular target. Machine-learning models can then learn structure– activity relationships from these screening data and prioritize compounds from larger libraries, focusing experimental time and resources on a smaller subset. This approach can be applied to collections such as the Drug Repurposing Hub (DRH) [6], whose compounds are chemically and pharmacologically annotated and have existing development information that can inform experimental follow-up.

Graph Neural Networks (GNNs) learn these relationships directly from molecular graphs, with atoms represented as nodes and bonds as edges, using message passing to combine information from local neighborhoods [7, 8]. Within the GNN family, a Directed Message-Passing Neural Network (D-MPNN) [9], implemented in Chemprop [10] and trained on an *Escherichia coli* screen, led to the identification of halicin [11]. The same general approach was subsequently applied to an *A. baumannii* screen, leading to the identification of abaucin through experimental follow-up [12].

Alongside prioritizing compounds, understanding which molecular features support their predicted activity can help formulate chemical hypotheses for experimental follow-up. Explainable artificial intelligence (XAI) provides tools for examining what a predictive model has learned and can support knowledge extraction [13] when the learned relationships reflect relevant aspects of antibacterial activity [14]. For the abaucin D-MPNN, such an interpretation must account for both its learned graph representation and the roughly 200 normalized RDKit descriptors supplying additional molecule-level physicochemical information [10, 15]. Removing these descriptors reduced predictive performance in the abaucin D-MPNN [12], consistent with benefits reported for other graph-based models [16, 17], although their contribution is dataset-dependent [18]. We therefore ask whether a graph-only model can retain comparable predictive performance while allowing atom and substructure contributions to be assessed without a separate molecule-level descriptor input.

In antibacterial discovery, Wong et al. [19] identified connected substructures that retain high predicted activity when evaluated on their own, then grouped these rationales to guide the discovery of structural classes of antibiotics. More generally, attribution methods express the importance assigned to molecular features as scores. Benchmark studies have evaluated such scores against known relevant features in tasks with computable ground-truth explanations [20]. The correspondence between these scores and the model’s behavior can be assessed by examining how predictions respond when highly attributed features are perturbed and whether the scores depend on the learned model parameters [21, 22]. However, even faithful attributions describe the behavior of the predictive model rather than establishing a biological mechanism [14].

Complementing the search for subgraphs that retain predicted activity, we assess recurring motifs by perturbing their constituent atoms within each parent molecule and comparing the resulting changes in predicted activity between experimentally active and inactive molecules containing the motif. However, the magnitude of a decrease is constrained by the original prediction: a probability already close to zero has less scope to decrease than one close to one. When comparing molecules that contain the same motif, differences between experimentally labeled active and inactive molecules can therefore reflect their initial predicted probabilities rather than a specific reliance on the motif.

We therefore compare the motif’s perturbation effect with the effects of perturbing equally sized sets of randomly selected atoms in the same molecule before contrasting its contribution between active and inactive molecules.

Prospective screening must ultimately translate model predictions into compounds selected for experimental testing. Yet prediction scores alone do not account for structural novelty, developability or potential toxicity, all of which inform the selection of compounds for follow-up. This motivates a prespecified ranking and triage procedure that combines predicted activity with these additional criteria.

Here, we investigate whether graph-only prediction can support antibacterial repurposing against *A. baumannii* through both chemical interpretation and prospective candidate selection (Figure 1). We assess predictive performance against the descriptor-augmented D-MPNN of Liu et al. [12] and evaluate whether the explanations reflect model behavior. Extending this evaluation from individual atoms to recurring substructures, we ask which motifs contribute more strongly to predictions in experimentally active than inactive molecules when their effects are assessed relative to other perturbations within the same molecule. In parallel, we apply the predictor to the Drug Repurposing Hub, with candidates prioritized using predicted activity, structural novelty, developability, predicted toxicity [23] and structural alerts [24, 25]. From this shortlist, CE3F4 is taken to a preliminary *in vitro* assay.

**Figure 1:**
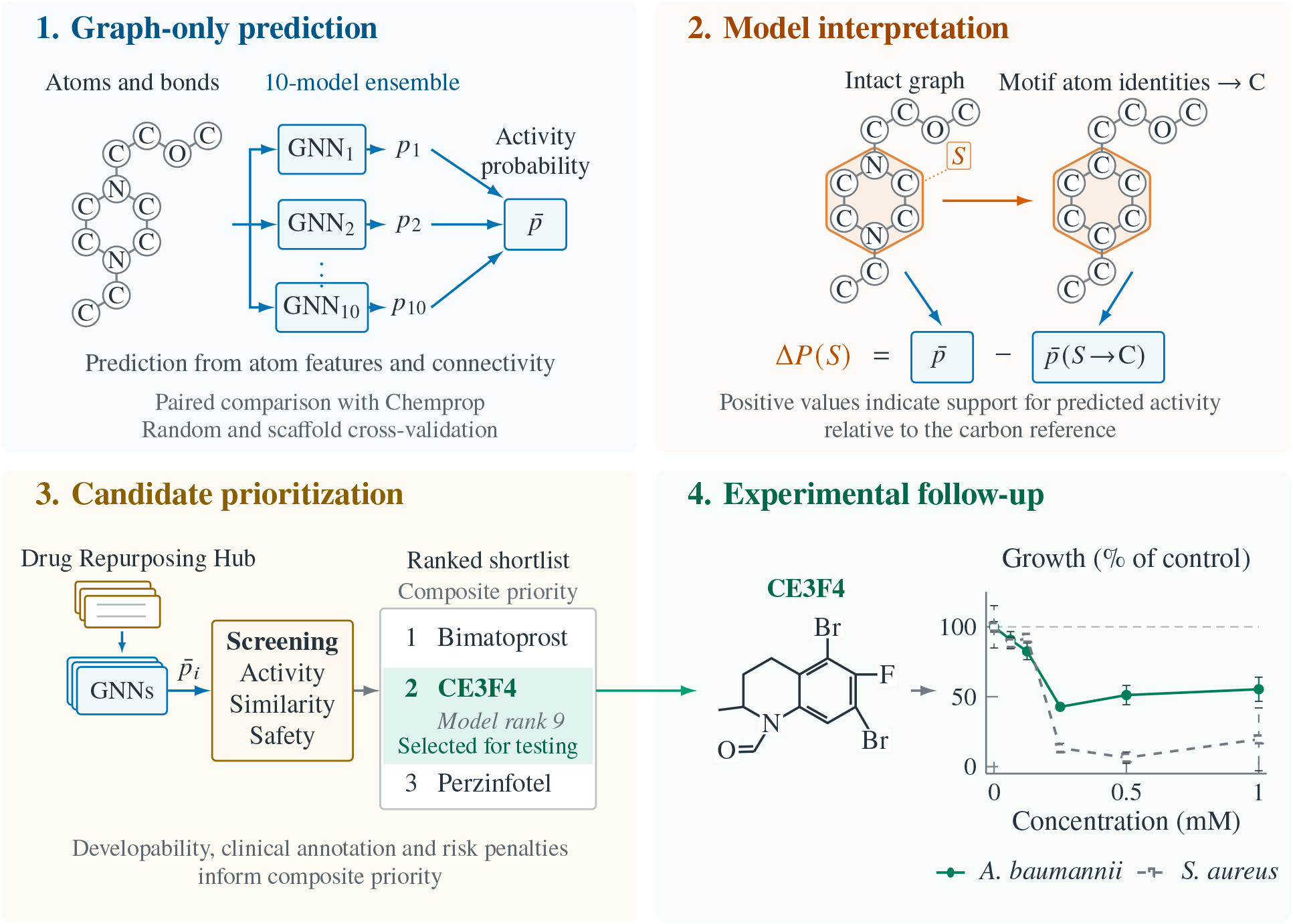
Graph-only prediction, evaluation of chemical explanations and prospective testing. **(1)** Ten independently initialized GNNs predict antibacterial activity from molecular graphs, and their probabilities are averaged to obtain 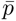. **(2)** Joint motif occlusion replaces the element identities of the shaded atoms with carbon, retaining all other features and connectivity. The same ensemble evaluates both inputs, and their probability difference defines the motif attribution Δ*P* (*S*). A positive value indicates that the original atom identities support predicted activity relative to the carbon reference. Attributions are evaluated through perturbation and restoration, parameter randomization assessed through occlusion, and agreement with Integrated Gradients. Motif effects are also compared with equally sized random atom sets within each molecule before contrasting experimentally active and inactive motif carriers. **(3)** Ensemble predictions support activity and similarity screening of the Drug Repurposing Hub, followed by safety assessment and composite prioritization incorporating developability, clinical annotation and risk penalties. CE3F4 ranks ninth by predicted activity among filter-passing candidates and second by composite priority, and was selected for testing. **(4)** The racemic CE3F4 structure is shown alongside measured growth relative to matched vehicle controls after 16 h. Points and error bars show technical-replicate means and SD from Data S4; the dashed line marks control growth. Molecular graphs and library records are schematic; rankings and assay measurements are study results.

## 2 Materials and Methods

### 2.1 Datasets and Data Curation

Following Liu et al. [12], we use their primary screen of 7 684 molecules tested for growth inhibition of *A. baumannii in vitro* as our training set. Following the original activity definition, we label a molecule as active when its growth value is below the dataset mean minus one standard deviation, resulting in 480 active and 7 204 inactive molecules (a *∼* 6.2% positive rate). Because the original screen tested each compound at 50 *µ*M [12], the activity labels and the predictions derived from them refer to that assay concentration.

For prospective screening, we use the 2020 release of the Drug Repurposing Hub [6]. Before graph construction, both datasets are deduplicated by canonical SMILES [26] (first occurrence kept), and entries that fail to parse or fall outside the range of heavy atoms [2, 200] are discarded. These filters do not remove molecules from the training screen (all 7 684 parse and are unique) but reduce the DRH from its 6 680 compounds to 6 642. Multi-fragment training entries such as salts and mixtures are retained as provided.

### 2.2 Molecular Graph Construction

We convert each molecule from its SMILES representation to a molecular graph *G* = (*V, E*) with RDKit [15], where nodes are atoms and edges are bonds. We use the same implicit-hydrogen treatment as the Chemprop baseline, avoiding graph expansion with additional hydrogen nodes. This also keeps prediction and attribution analyses on the same graph representation. Every node *v* carries a feature vector **x***_v_ ∈* ℝ^34^ combining a one-hot encoding of atom type in *{*0, 1*}*^22^, a one-hot encoding of total atom degree in *{*0, 1*}*^8^, and four structural features describing aromaticity, ring membership, radical-electron count and chirality. The degree encoding includes hydrogens even though they are not added as graph nodes. The atom-type encoding distinguishes {H, B, C, N, O, F, Na, Mg, Si, P, S, Cl, K, Ca, Fe, Cu, Zn, As, Se, Br, I, other}. In addition to the atom features, we compute bond feature vectors **e***_uv_ ∈* ℝ^7^ for all molecules, containing a one-hot encoding of bond type together with indicators of conjugation, ring membership and stereochemistry. These bond attributes are incorporated into message passing only by architectures that use edge features in their updates.

### 2.3 Model Architecture and Training

We selected the message-passing backbone, graph-level readout and prediction head through an Optuna search [27]. The candidate backbones included a Graph Isomorphism Network (GIN) [28], its bond-feature variant GINE [29], a Graph Attention Network (GAT) [30, 31] and the GATv2 variant [32]. Alongside these architecture choices, the search evaluated graph-level readouts based on averaging, summation or componentwise maximization of atom representations. A further readout concatenated the vectors obtained by global mean and max pooling, allowing the prediction head to use both average activations and per-feature maxima. Head depth and width, dropout and the use of residual connections were varied jointly with these backbone and readout choices. To reduce dependence on a single data split, the strongest configurations were re-evaluated under the full cross-validation protocol of Section 2.4. The full search space and configuration ranking are provided in Supporting Information S1.

The selected GIN backbone predicts antibacterial activity from atom features and graph connectivity, without incorporating bond feature vectors or molecule-level descriptors. To obtain this prediction, the model updates atom representations through *K* message-passing layers, combines them into a graph-level embedding, and passes this embedding to a multilayer perceptron (MLP) prediction head. To initialize the atom representations, each node feature vector **x***_v_* is projected into a *d*-dimensional space, 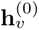 = ReLU(**W**_proj_**x***_v_* + **b**_proj_).

At layer *k*, the GIN updates node *v* by combining its own representation with the sum of the representations of its neighbors *N* (*v*) and passing the result through an MLP:

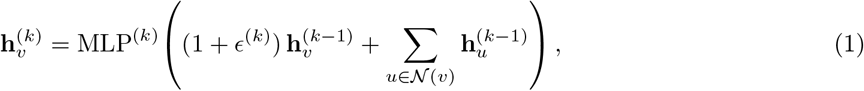

where *ɛ*^(*k*)^ is a learnable scalar and MLP^(*k*)^ is a two-layer perceptron with internal batch normalization. Each GIN layer’s output then passes through a ReLU and dropout. In the readout phase, sum pooling over the final node states, 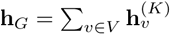, gives a graph-level embedding that an *L*-layer MLP maps to a scalar logit 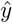. The corresponding probability is 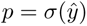.

During model selection, we compared unweighted binary cross-entropy (BCE), class-weighted BCE and focal loss [33] to assess alternatives for handling the class imbalance. For class-weighted BCE, the positive-class weight was the ratio *N^−^/N* ^+^ of inactive to active molecules in the training partition. The selected configuration uses unweighted BCE, so each model minimizes

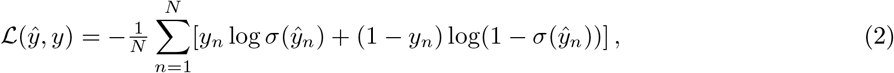

where *N* is the number of training molecules and *y_n_ ∈ {*0, 1*}* is the activity label. Optimization uses Adam with a weight decay of 1.32 *×* 10*^−^*^6^ and batches of 32 molecules. The learning-rate schedule first increases the rate to a maximum of 3.98 *×* 10*^−^*^4^ and then progressively decreases it. During model selection and cross-validation, early stopping monitors ROC-AUC on a held-out validation partition. For cross-validation, we retain the model parameters with the highest validation ROC-AUC.

The selected architecture uses *K* = 4 message-passing layers with hidden dimension *d* = 256 and sum pooling. The three-layer prediction head has 256 hidden units and dropout 0.262. Residual connections and additional normalization between GIN layers are omitted, while the internal batch normalization of each GIN layer is retained. With this configuration, the deployed predictor averages the probabilities of *M* = 10 independently initialized GIN models, 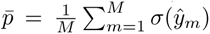. For prospective screening, this ensemble is trained on all labeled data for up to 256 epochs. Both predictions and attribution analyses use this ensemble.

### 2.4 Cross-Validation and Baseline Comparison

We evaluate predictive performance using complementary random and scaffold-based 10-fold cross-validation protocols. The random protocol follows the evaluation approach of Liu et al. [12], assigning molecules to folds without separating their scaffolds. The scaffold protocol instead keeps molecules sharing a Bemis– Murcko scaffold [34] in the same fold, assessing generalization to molecules whose scaffolds are absent from the corresponding training partition. In both protocols, predictions are averaged over a 10-model ensemble per fold, and metrics are computed on the held-out test partition. The exact fold construction is detailed in Supporting Information S2. We report Receiver Operating Characteristic Area Under the Curve (ROCAUC) and Precision-Recall Area Under the Curve (PR-AUC), the latter being particularly informative given the class imbalance.

Our baseline is the Chemprop D-MPNN in the configuration published by Liu et al. [12]. This model supplements its learned graph representation with approximately 200 normalized molecular descriptors computed with RDKit, summarizing physicochemical properties, molecular topology and functional-group counts. These include molecular weight, estimated lipophilicity, topological polar surface area and hydrogenbond donor and acceptor counts. The descriptor vector is concatenated with the learned graph representation before the final prediction. We use the authors’ default D-MPNN hyperparameters, with 3 message-passing steps, hidden size 300, two feed-forward layers, dropout 0 and 30 training epochs, and average predictions over a 10-model ensemble. The baseline’s random-split ROC-AUC of 0.797 is consistent with the 0.792 reported in the reference study. Chemprop is trained and scored on the same fold assignments as the GIN for both split protocols, making the comparison paired.

Because both models are evaluated on the same held-out molecules in each fold, we analyze their per-fold performance differences with paired *t*-tests and Wilcoxon signed-rank tests [35], reporting 95% Student-*t* confidence intervals for the mean difference. This pairing accounts for differences in test-fold difficulty by comparing the two models within each fold. We apply Holm correction to the four paired *t*-tests covering ROC-AUC and PR-AUC under both split protocols. Since a non-significant difference does not establish equivalence, we also use two one-sided tests (TOST) [36] to assess whether the mean paired difference lies within a specified margin of *±*0.05 AUC units. However, the folds share training data, so the uncertainty estimates and tests, including TOST, do not fully account for dependence between folds and are interpreted as approximate [37].

### 2.5 Prospective Screening and Candidate Prioritization

The deployed GIN ensemble, trained on all labeled data, assigns each Drug Repurposing Hub compound an averaged predicted probability of antibacterial activity, 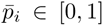, using the graph representation of Section 2.2. Following the screening criteria of Liu et al. [12], we retain compounds with 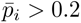 and a maximum Tanimoto similarity below 0.3 to any experimentally active training molecule. Similarity is computed using Morgan fingerprints [38] with radius 2 and 2048 bits. This similarity quantifies structural dissimilarity from known active compounds, while scaffold-based cross-validation (Section 2.4) assesses generalization to unseen scaffolds.

Candidates passing these filters are then assessed for developability and potential safety liabilities using physicochemical properties, clinical annotation, predicted toxicity and structural alerts. Compounds whose canonical SMILES match a training molecule exactly are flagged, and the assessment retains these matches through scoring. The prospective subset excludes these matches, while their presence in the scored table allows the effect of this exclusion on the shortlist to be checked. Supporting Information S5 distinguishes the filter-passing, annotated and prospectively eligible populations.

#### Annotation

Each compound is annotated against the Broad Drug Repurposing Hub [6] to recover its identity, clinical status, mechanism-of-action class, and a known-antibiotic flag, so that rediscoveries of existing antibacterials are identified rather than counted as novel hits.

#### Predicted toxicity and structural alerts

We predict six toxicity-relevant endpoints with ADMETAI [23]: hERG inhibition, clinical toxicity (ClinTox), drug-induced liver injury (DILI), acute toxicity (LD_50_), Ames mutagenicity and carcinogenicity. Compounds are also screened against a curated set of structural alerts comprising 15 reactive-chemistry patterns and 18 nuisance or assay-risk patterns. These alerts are encoded as SMARTS [39] queries and simple property-based heuristics and are listed in the Supporting Information (Section S5). They were informed by the literature on reactive and nuisance substructures, but do not correspond to the complete published PAINS [24] and Brenk [25] catalogues, which were not applied at this stage. The single compound selected for experimental testing was subsequently screened against the complete catalogues, as reported in Section 3.

#### Prioritization

A heuristic composite priority score combines predicted activity with structural dissimilarity, physicochemical properties and clinical annotation to produce a ranked shortlist for experimental testing. Predicted toxicity, available safety information and structural alerts contribute penalties to the ranking. These criteria overlap: structural novelty contributes through both continuous similarity and a binary novelty indicator, while safety annotation informs both translational priority and risk penalties. They are therefore treated as weighted prioritization criteria rather than independent measures of candidate quality. This *in silico* triage aims to prioritize compounds with a favorable predicted safety profile, but does not replace experimental cytotoxicity or selectivity measurements, which remain necessary follow-up work. Score components, column definitions and thresholds are listed in Supporting Information S5.

### 2.6 Interpretability Analysis

We interpret the deployed GIN ensemble using atom-level attribution scores, which describe each atom’s contribution to a specified model output relative to a reference input. We use two complementary methods, atom occlusion and Integrated Gradients (IG) [40], with carbon as the reference atom type. The reference operation replaces an atom’s element-identity encoding with that of carbon while retaining the other atom features, atom count and connectivity. This probes the contribution of element identity while holding graph connectivity and the other atom features fixed. Zeroing an atom’s feature vector would instead leave a “ghost atom”: a connected node whose encoding specifies no atom type.

Atom occlusion, adapted from occlusion sensitivity [41], measures the change in the ensemble’s predicted probability when one atom’s identity is replaced with carbon, 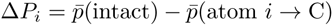. A positive value indicates that the atom’s original identity supports the prediction of activity relative to the carbon reference. IG is computed on each GIN model’s logit using Captum [42], with 128 integration steps along a path in feature space from a reference graph with all atom identities set to carbon to the original graph representation. The feature attributions are averaged across the ensemble and summed over feature dimensions to obtain a score for each atom.

#### Faithfulness

We assess faithfulness by testing whether perturbing atoms ranked as important changes the prediction more than perturbing randomly ordered atoms, using deletion and insertion curves [21]. For deletion, we start from the original graph and progressively replace atom identities with carbon in decreasing attribution order. Conversely, insertion starts from the reference graph in which all atom identities are set to carbon and progressively restores their original identities in the same order. We track the ensemble’s predicted probability 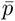 as a function of the fraction of atoms perturbed or restored and compare the area under each curve with that obtained under a random ordering. The per-molecule differences in area are assessed with one-sided paired Wilcoxon signed-rank tests. Lower deletion AUC and higher insertion AUC than under random ordering support the attribution ranking under these perturbations.

We evaluate faithfulness on all 480 training actives and on the top-scoring Drug Repurposing Hub molecules absent from the training set, after removing exact canonical-SMILES matches, to assess whether attribution rankings remain informative on unseen molecules.

#### Model-randomization test

Following Adebayo et al. [22], we assess the dependence of occlusion attributions on the learned parameters by comparing the trained GIN ensemble with ten independently reinitialized ensembles. For each molecule, we compute the Spearman correlation between trained and randomized atom-level attributions. We summarize each randomization by its median correlation across molecules and report the mean and standard deviation of these medians over the ten randomizations. As a control for consistency within the trained ensemble, we compare attributions obtained from two disjoint groups of five trained GIN models and report the median correlation across molecules.

#### Inter-method agreement

We compare the per-atom rankings produced by occlusion and IG using Spearman correlation within each molecule, reporting the median correlation across molecules as a measure of agreement between the methods.

#### Data-driven substructure enrichment

From occlusion attributions on active molecules, we extract the most-attributed atoms and their ring systems as candidate fragments, deduplicating stereoisomeric variants. For each recurring fragment, we jointly perturb its constituent atoms using the same carbon-substitution operation and compare the resulting change in predicted probability between experimentally active and inactive molecules containing the fragment. We test this difference with a one-sided Mann–Whitney *U* test [43] and control the false-discovery rate (FDR) using the Benjamini–Hochberg procedure [44], with significance defined by *q <* 0.05.

#### Baseline-controlled enrichment

Because the predicted probability after perturbation cannot fall below zero, the raw decrease Δ*P* cannot exceed the model’s original predicted probability for the intact molecule. A molecule assigned a low probability of activity therefore has less scope for a large decrease after perturbation, making raw effects difficult to compare across molecules with different initial scores. We therefore report a within-molecule control for the perturbation effect. For a substructure occupying the atom set *S* with 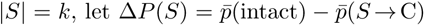 denote the change in predicted probability when its constituent atoms are perturbed together. We draw *R* = 3 random atom subsets *S_r_* of the same size *k* from the atoms of the same molecule and define the excess importance:

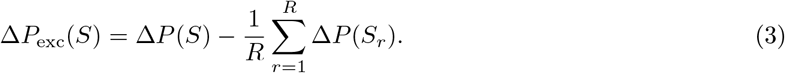

This comparison holds the molecule and the number of sampled atoms fixed, so a positive Δ*P*_exc_ means that perturbing the substructure decreases the predicted probability more than perturbing an equally sized set of atoms drawn from the same molecule. Carbonization rewrites only the element-identity encoding, so it leaves an atom that is already carbon unchanged: the control matches the number of atoms sampled, not the number whose encoding actually changes. Measured over the carriers, this asymmetry is small for piperazine (2.0 non-carbon atoms in the motif against 1.8 in the equal-size samples) and favours the control for four of the six motifs examined. Therefore, the comparison is conservative rather than permissive for those motifs. An all-carbon motif is unchanged by the operation by construction, which is why benzene shows a negative excess.

We then compare this excess importance between active and inactive carriers using the same one-sided Mann–Whitney *U* test and Benjamini–Hochberg correction, with *q <* 0.05. A significant enrichment supports greater relative reliance on the motif in experimentally active molecules under this control. The sensitivity of this control to the number of random atom sets and to the choice of null is examined in Supporting Information S4.

### 2.7 *In vitro* Assay

We measured growth inhibition of CE3F4 against *A. baumannii* NCTC 13305 (assembly GCA_900444775) and, as a Gram-positive non-target comparator, *S. aureus* M/8912 (assembly CABEFO010000000), using broth micro-dilution. NCTC 13305 is a clinical isolate distributed as a reference control for OXA-58-like carbapenemase detection and is distinct from the strain used to generate the training labels, *A. baumannii* ATCC 17978. A 20 mM stock of CE3F4 in DMSO was prepared and serially diluted two-fold. Precultures were grown in tryptic soy broth (TSB) for 8 h at 37*^◦^*C with shaking at 200 rpm, then diluted to OD_600_ = 0.05. Each well of a 96-well plate received 95 *µ*L of culture and 5 *µ*L of compound solution, giving final CE3F4 concentrations of 0.0625–1 mM and 5% (v/v) DMSO. The compound-free control contained the same DMSO concentration, as higher DMSO concentrations themselves inhibited growth. After incubation for 16 h at 37*^◦^*C and 200 rpm, OD_600_ was measured on a Tecan Spark. Relative growth was calculated as the blank-corrected OD_600_ expressed as a percentage of the 5% DMSO control. Each condition was measured in at least three technical replicate wells dispensed from a single preculture per strain. These wells therefore quantify within-assay variability rather than variation between independent biological cultures. Racemic CE3F4 (CAS 143703-25-7) was purchased from Szabo-Scandic, catalogue MEXHY-108539-10, supplied as MedChemExpress HY-108539, lot 279781. The supplier’s Certificate of Analysis (COA) for this lot reports 98.85% purity by HPLC, with ^1^H NMR and MS consistent with the structure, and it is available in Supporting Information S7.

Because CE3F4 is poorly water-soluble and its enantiomer CE3F4R has a reported critical aggregation concentration near 150 *µ*M [45], well within our assayed range, aggregation remains a possibility. Although no precipitate was visible between 0.0625 and 1 mM, visual inspection cannot rule out sub-visible colloids. We therefore report concentrations as nominal rather than confirmed free concentrations. No detergent-based aggregation counter-screen was performed [46].

#### Safety

Both tested strains, *A. baumannii* NCTC 13305 and *S. aureus* M/8912, are human pathogens. All handling of live cultures was performed at Austrian Institute of Technology (AIT) in a biosafety cabinet within a Biosafety Level 2 (BSL-2) laboratory. Dimethyl sulfoxide readily crosses skin and carries dissolved solutes with it, so standard personal protective equipment (gloves, lab coat and eye protection) was used when handling compound stocks.

## 3 Results and Discussion

We first assess whether the graph-only model performs comparably to the descriptor-augmented baseline. We then examine whether its chemical explanations reflect model behavior and how its predictions support candidate selection for experimental follow-up.

### 3.1 Classification Performance

We compare the selected GIN configuration with the reference Chemprop D-MPNN using fold-specific ensembles trained and evaluated on identical partitions (Table 1).

**Table 1:** Predictive performance of the graph-only GIN and descriptor-augmented Chemprop D-MPNN [12] under random and scaffold-based cross-validation. Both methods use identical partitions and ten-model ensembles within each fold. Values are mean *±* SD across the ten folds. Δ denotes GIN minus Chemprop, and counts in parentheses indicate folds with higher GIN performance. The two *p*-values correspond to paired *t*-tests and Wilcoxon signed-rank tests, respectively. The 95% confidence intervals are based on the Student-*t* distribution and refer to the mean paired difference.

| Split | Metric | GIN | Chemprop | $\Delta$ (wins) | $t$ / Wilcoxon $p$ | $\Delta$ 95% CI |
| --- | --- | --- | --- | --- | --- | --- |
| Random | ROC-AUC | $0.819 \pm 0.031$ | $0.797 \pm 0.038$ | +0.022 (9/10) | 0.006 / 0.004 | [+0.008, +0.035] |
| Random | PR-AUC | $0.370 \pm 0.060$ | $0.374 \pm 0.062$ | -0.004 (3/10) | 0.73 / 0.43 | [-0.030, +0.021] |
| Scaffold | ROC-AUC | $0.783 \pm 0.046$ | $0.761 \pm 0.046$ | +0.022 (9/10) | 0.09 / 0.06 | [-0.004, +0.048] |
| Scaffold | PR-AUC | $0.311 \pm 0.080$ | $0.303 \pm 0.090$ | +0.008 (4/10) | 0.77 / 1.0 | [-0.053, +0.069] |

Under random splitting, the GIN achieves a small ROC-AUC advantage over Chemprop, 0.819 versus 0.797, with a mean paired difference of +0.022 and a Holm-adjusted *p* = 0.025 across the four benchmark comparisons. This advantage does not extend to PR-AUC, for which the respective means are 0.370 and 0.374. Under scaffold splitting, predictive performance decreases for both models, and the mean ROC-AUC difference remains +0.022, but its 95% confidence interval includes zero. The corresponding PR-AUC means are 0.311 and 0.303, with no statistically detectable difference.

To distinguish an undetected difference from evidence of equivalence, we also examine the TOST results within the specified *±*0.05 margin. These support equivalence for random-split ROC-AUC and PR-AUC and for scaffold-split ROC-AUC, whereas scaffold-split PR-AUC remains inconclusive. As with the difference tests, these results are approximate because the cross-validation folds share training data.

Taken together, these results show that the graph-only model retains competitive predictive performance without the additional molecular descriptors used by the baseline. The comparison includes a small randomsplit ROC-AUC gain, while equivalence for scaffold-split PR-AUC remains unresolved. Trial 210 was selected for its highest cross-validated ROC-AUC among the five re-evaluated configurations, so the same evaluation results served both model selection and performance reporting. The GIN was tuned over 208 completed Optuna trials, whereas Chemprop used its published configuration, so the comparison concerns the two modeling workflows rather than an isolated change in architecture or descriptors. Configuration rankings and per-fold results for both models and split protocols are provided in Supporting Information S1 and S2, respectively.

### 3.2 Model Interpretation

Having assessed predictive performance, we next examine whether the model’s explanations identify chemical features that support its predictions. We first evaluate the attribution rankings against the model’s responses to perturbations, then investigate recurring substructures across active and inactive molecules.

#### Attribution faithfulness and model dependence

Figure 2 shows that replacing atom identities with carbon in decreasing attribution order reduces the predicted activity probability faster than a random ordering. Conversely, restoring the original atom identities in the same order recovers the probability faster. These responses indicate that occlusion and Integrated Gradients prioritize atoms whose element identities support the model’s prediction of activity relative to the carbon reference. We summarize these trajectories by the area under each probability curve across the fraction of atoms perturbed or restored. Lower deletion area and higher insertion area therefore indicate more effective attribution rankings. Across the 480 training actives, mean deletion areas are 0.10 for occlusion and 0.09 for Integrated Gradients, compared with 0.19 for random ordering. Mean insertion areas are 0.31 and 0.32, respectively, compared with 0.20 for random ordering. The attribution-guided ordering outperforms random ordering in 92–98% of molecules across the two attribution methods and curve types, with paired Wilcoxon signed-rank *p <* 10*^−^*^10^ in each comparison.

**Figure 2:**
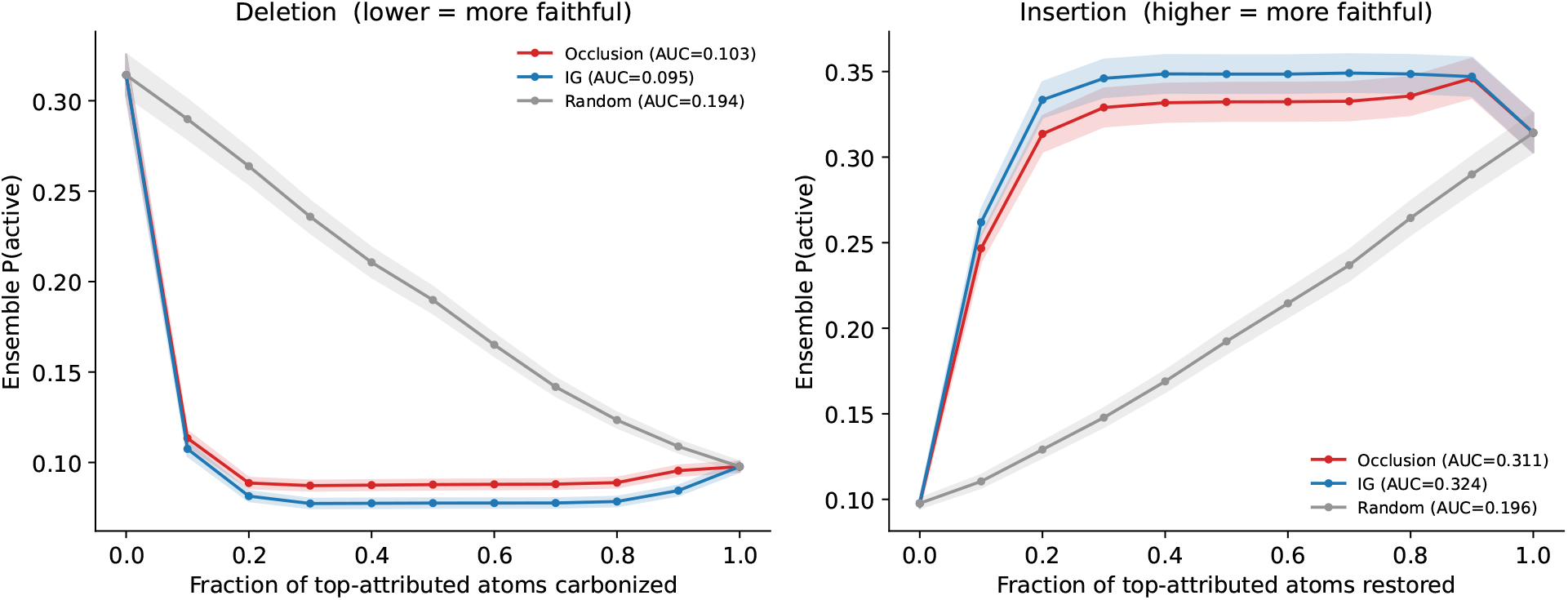
Faithfulness of occlusion and Integrated Gradients on the 480 training actives. Deletion (left) progressively replaces atom identities with carbon in decreasing attribution order. Insertion (right) starts with all atom identities set to carbon and progressively restores their original identities in the same order. The rankings from occlusion (red) and Integrated Gradients (blue) are compared with a random ordering (grey). Other atom features and graph connectivity remain fixed. Curves show the mean predicted activity probability of the GIN ensemble across molecules, with shaded bands representing one standard error of the mean. The AUC values in the legends are areas under these probability curves over the fraction of atoms perturbed or restored. Lower deletion area and higher insertion area than under random ordering indicate that the attributions prioritize atom identities supporting predicted activity relative to the carbon reference.

This separation also holds on the top-scoring Drug Repurposing Hub molecules absent from the training set, as shown in Supporting Information Figure S1. Mean deletion areas are 0.12 for occlusion and 0.11 for Integrated Gradients, compared with 0.25 for random ordering. Occlusion and Integrated Gradients each yield a mean insertion area of 0.44, compared with 0.25 for random ordering. The attribution-guided ordering outperforms random ordering in 97–99% of these molecules, with *p <* 10*^−^*^10^ in each paired comparison. These results extend the evidence for faithfulness to prospective-library molecules not used for model training.

For occlusion, the attribution calculation and the evaluation curves use the same carbon-substitution operation, so their agreement establishes consistency under that perturbation. Integrated Gradients derives its ranking from gradients and also outperforms random ordering in these probability-based evaluations, providing complementary support for faithfulness.

To assess whether the explanations depend on what the model has learned, Figure 3 compares occlusion attributions before and after parameter randomization. Across ten randomizations, the median per-molecule

**Figure 3:**
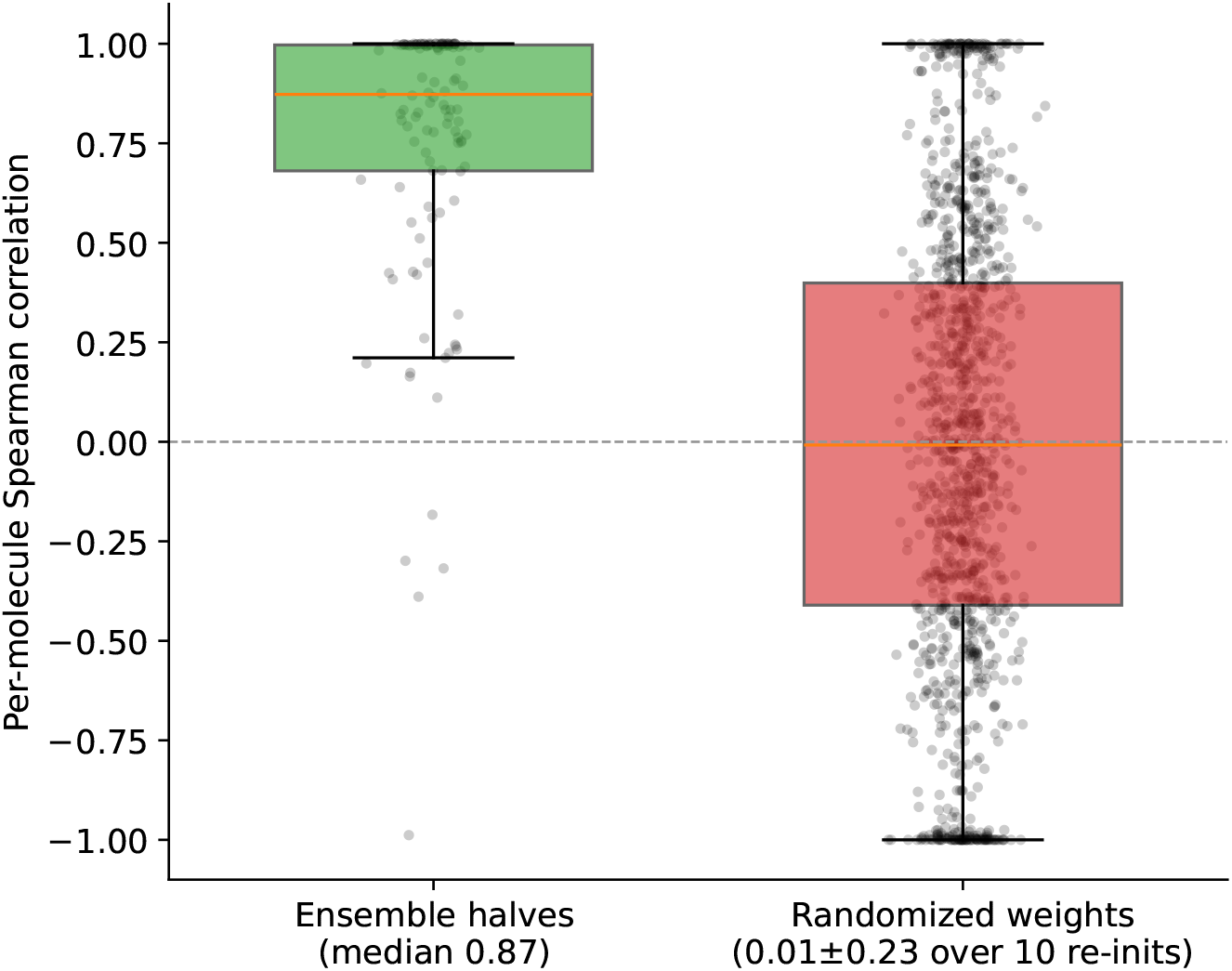
Dependence of occlusion attributions on learned model parameters. The green distribution shows per-molecule Spearman correlations between attributions from two disjoint groups of five trained GIN models, with median 0.87. The red distribution shows correlations between attributions from the trained ensemble and ten independently reinitialized ensembles, pooled across molecules and randomizations. Points denote individual correlations, and boxes show medians and interquartile ranges. The 0.01 *±* 0.23 annotation reports the mean and standard deviation of the ten per-randomization median correlations. The lower agreement after randomization supports dependence on learned model parameters.

Spearman correlations have a mean of 0.01 and a standard deviation of 0.23, whereas the median correlation between two disjoint halves of the trained ensemble is 0.87. This separation supports dependence on the learned parameters. Occlusion and Integrated Gradients also agree on atom rankings, with a median per-molecule Spearman correlation of 0.72, as shown in Supporting Information Figure S2. Together, the perturbation tests, parameter randomization and inter-method agreement support examining the chemical features highlighted by the attributions.

#### Substructure enrichment and within-molecule control

We then move from individual atom rankings to recurring substructures, testing whether perturbing a given motif produces a larger decrease in the model’s predicted activity probability in experimentally active than inactive molecules containing it. Figure 4, left, shows five motifs with greater raw importance in active molecules: a bicyclic thioether core, piperazine, 4-aminotetrahydropyran, 2-aminothiazole and pyridine. Each is significant under a one-sided Mann–Whitney *U* test with Benjamini–Hochberg correction at *q <* 0.05.

**Figure 4:**
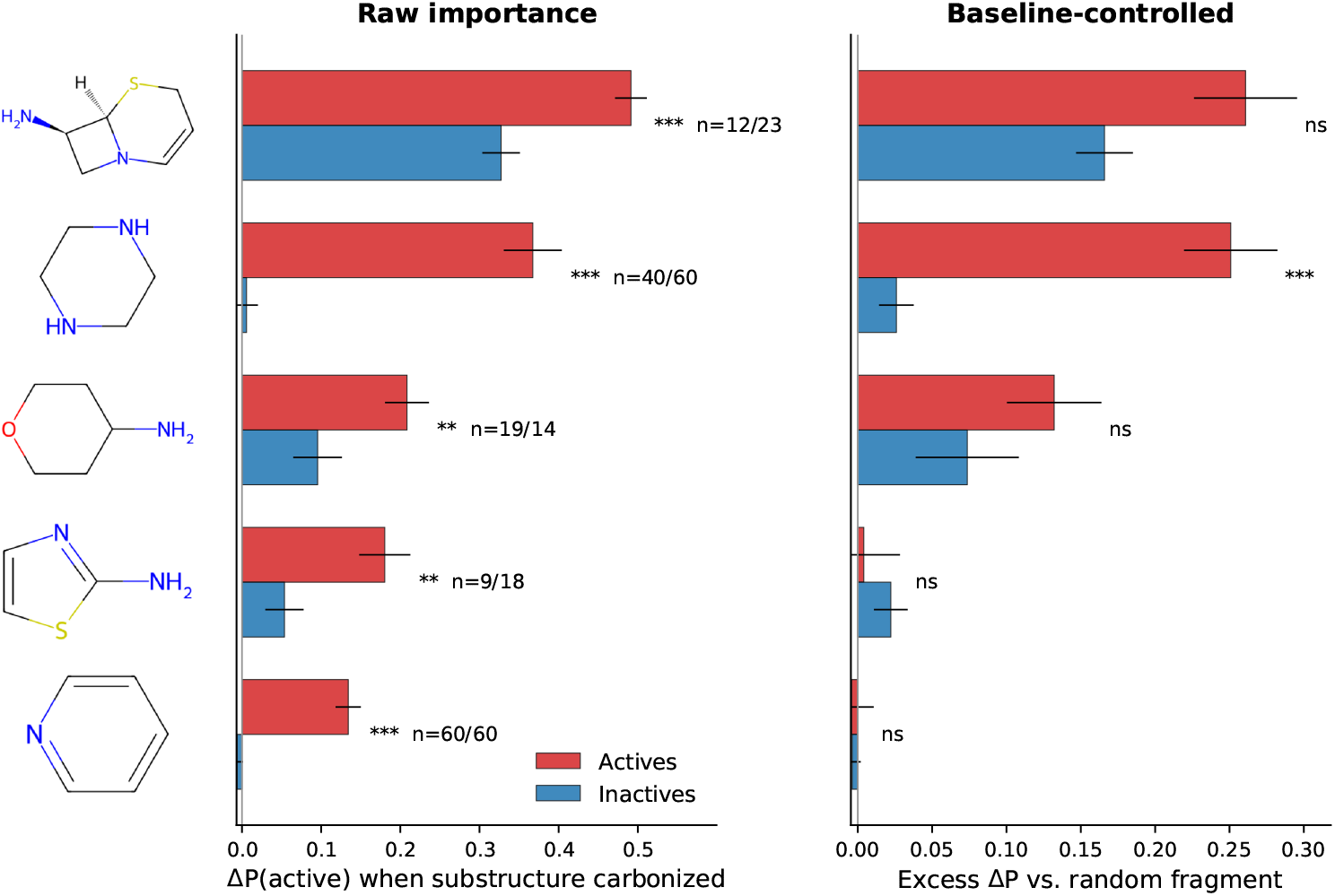
Substructure importance in experimentally active and inactive molecules. Bars show mean raw importance Δ*P* (left) and within-molecule excess importance Δ*P*_exc_ (right), for active (red) and inactive (blue) molecules containing each motif. Raw importance measures the decrease in the ensemble’s predicted activity probability after carbon substitution of the motif’s atom identities. Excess importance subtracts the mean effect of *R* = 3 equally sized random atom sets within the same molecule. Error bars indicate standard errors of the mean across molecules containing the motif, computed separately for active and inactive groups. The counts labelled *n* give the numbers of active and inactive molecules, respectively, containing each motif. For each panel, significance is assessed using one-sided Mann–Whitney *U* tests with Benjamini–Hochberg correction. One, two and three stars denote *q* values below 0.05, 0.01 and 0.001, respectively, while *ns* denotes *q ≥* 0.05. Only piperazine remains significant after the within-molecule control.

Two of the enriched motifs correspond to parts of established antibiotic scaffolds. The bicyclic thioether motif matches the cephem skeleton of cefotaxime and ceftriaxone, rather than penicillins or carbapenems, but lacks the *β*-lactam carbonyl. Its attribution therefore concerns a portion of these antibiotic scaffolds that does not include that functional group. Similarly, in fluoroquinolones, the pyridine match lies within the 4-quinolone ring of the scaffold rather than in a pendant pyridine. Its co-occurrence with piperazine thus reflects the shared class skeleton and does not establish two independent pharmacophores.

However, a larger raw perturbation effect can also reflect a higher initial prediction, because a molecule with a low predicted probability has less room for that probability to decrease. We therefore compare each motif’s importance with the mean effect of equally sized random atom sets within the same molecule, as shown in Figure 4, right. In the primary analysis with *R* = 3, piperazine is the only significant motif after this control, with mean excess importance of Δ*P*_exc_ = 0.25 in active molecules versus 0.03 in inactive molecules and *q* = 7 *×* 10*^−^*^8^. The bicyclic thioether core is borderline in this analysis, with *q* = 0.058, while the other three motifs are not significant. Piperazine therefore provides the strongest evidence of a recurring chemical motif that supports activity predictions beyond the effect of equally sized random atom sets within the same molecule. Its basic, protonatable nitrogen atom occurs across antibacterial chemotypes, including fluoroquinolones, offering a chemical context for this model reliance. Amine-containing compounds have been associated with accumulation in Gram-negative bacteria [47], although those accumulation rules were derived in *E. coli* and emphasized primary amines. Their direct mechanistic transfer to piperazine-containing compounds in *A. baumannii* remains unconfirmed.

#### Robustness of the piperazine signal

To assess the robustness of the piperazine result, we first varied the number and connectivity of the random atom sets used in the within-molecule control. Piperazine remains strongly enriched when the number of sets increases from 3 to 100 or when sampling is restricted to connected subgraphs of equal size, with *q ≤* 4*×*10*^−^*^7^. We then separated fragment discovery from enrichment testing by nominating substructures on one half of the actives and testing them on the other half. Piperazine remains strongly enriched in this second group, with *q <* 10*^−^*^6^. Because the deployed ensemble was trained on all actives, this split checks the dependence on fragment selection rather than generalization of the trained model. Full sensitivity and split-half results, including weaker or control-dependent signals from other motifs, are reported in Supporting Information S4.

Finally, we examined the model’s reliance on these motifs in the Drug Repurposing Hub. Among piperazine-containing compounds, the change in predicted probability after perturbing piperazine correlates with the intact molecule’s score, with Spearman *ρ* = 0.84, and the association extends to the broader basic-amine class (Supporting Information S4). Together, the enrichment and library-wide analyses identify a recurring chemical feature used by the deployed model, providing a basis for experimental hypotheses about its role in antibacterial activity. Establishing a generalizable structure–activity relationship requires independent experimental validation.

Figure 5 illustrates this reliance on a representative active fluoroquinolone with an ensemble-predicted activity probability of 0.89. The two piperazine nitrogens receive the highest atom-level attributions, and perturbing the piperazine atoms together decreases the prediction by Δ*P* = 0.75. The corresponding deletion and insertion curves support the local faithfulness of this explanation.

**Figure 5:**
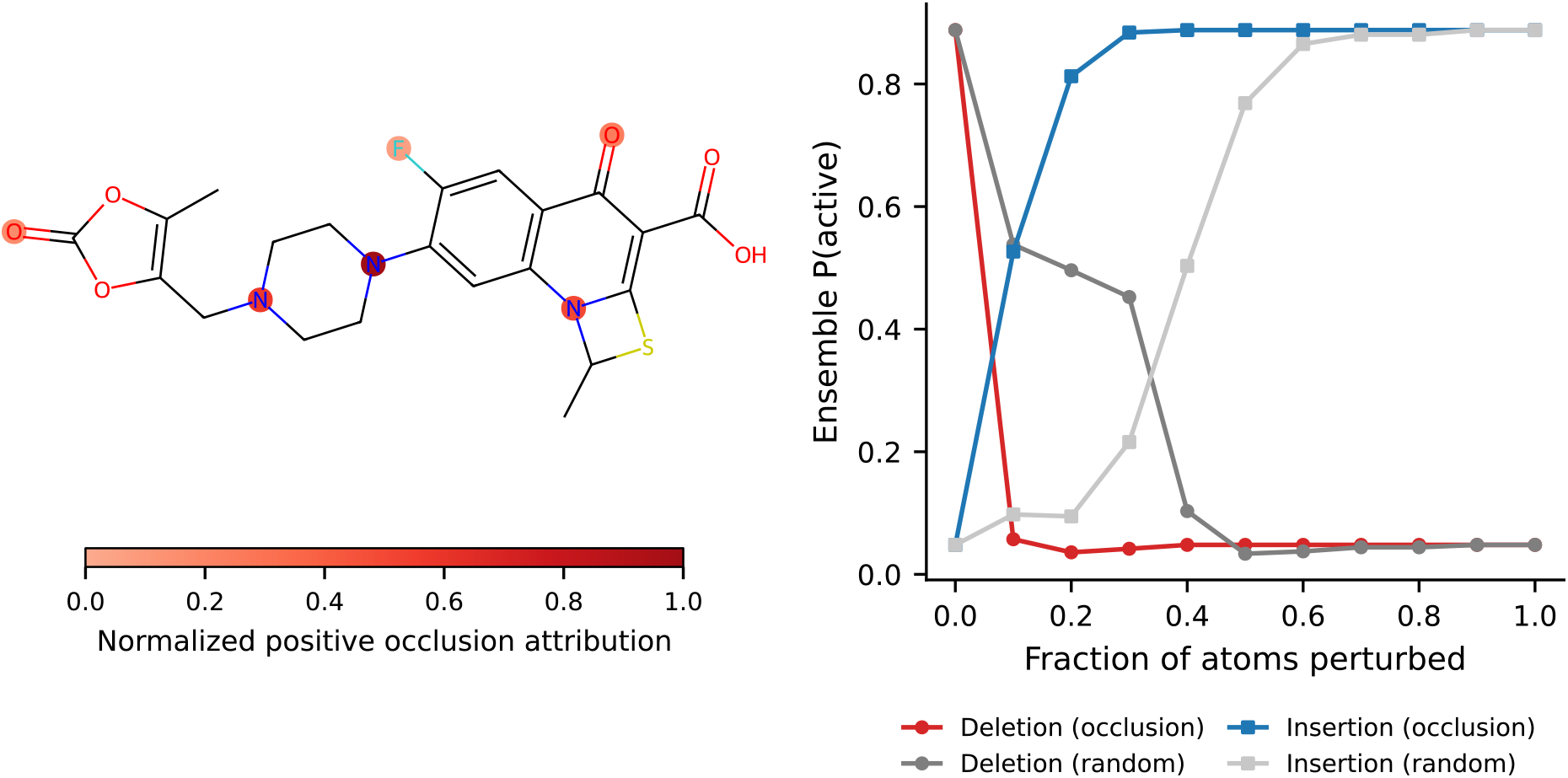
Atom-level explanation for an active, piperazine-containing fluoroquinolone. Left, red highlights show positive occlusion attributions scaled to the largest positive attribution in this molecule, with darker shading indicating greater importance. The piperazine nitrogens receive the highest scores. Perturbing the piperazine motif decreases the ensemble’s predicted activity probability by Δ*P* = 0.75 from its intact value of 0.89. Right, deletion and insertion curves compare the occlusion-guided ordering with a random ordering. Circles denote deletion and squares denote insertion, with colored curves for occlusion and grey curves for random ordering. The faster probability decrease and recovery support the local faithfulness of the occlusion ranking under carbon substitution.

### 3.3 Prospective Screen and Candidate Prioritization

The deployed GIN ensemble, trained on all labeled data, identified 418 Drug Repurposing Hub records meeting the screening criteria of Section 2.5, with predicted activity probability above 0.2 and maximum Tanimoto similarity below 0.3 to active training molecules. Twelve of these are exact matches to molecules in the training screen and two further records are duplicate entries for the same compound, so the 418 records correspond to 405 distinct candidates absent from the training screen. The annotation and scoring stage covers the 417 distinct compounds, including the twelve matches, none of which entered the shortlist.

Within this candidate pool, the highest-scoring compound is eravacycline [48], a marketed fluorocycline antibiotic, providing a qualitative check that the predictor assigns a high score to a known antibacterial. Its nearest active training neighbor is minocycline, with a maximum Tanimoto similarity of 0.25, and seven tetracycline-class compounds are present among the training actives. Its nearest neighbor in the screen as a whole is tigecycline, at 0.336, which the same activity rule labels inactive: the similarity criterion is defined against active training molecules only, following Liu et al. [12]. The high score assigned to eravacycline is therefore consistent with generalization from the tetracyclines among the training actives, rather than extrapolation beyond the training chemistry.

Eravacycline was retained as a qualitative positive reference and excluded from experimental prioritization as an existing antibiotic. Its passage through the similarity filter therefore illustrates why fingerprint dissimilarity must be supplemented with clinical annotation to distinguish repurposing candidates from existing antibiotics.

The subsequent developability and safety assessment produced the ranked shortlist in Table 2. The composite score combines activity predictions with structural dissimilarity, physicochemical properties, clinical annotation and risk penalties.

**Table 2:** Prioritized candidates for experimental follow-up, ordered by decreasing composite priority score. Model score 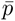 is the mean predicted activity probability of the GIN ensemble. *T*_max_ denotes the maximum fingerprint Tanimoto similarity to an active training molecule. Risk is the composite risk penalty, while Priority combines predicted activity, structural dissimilarity, physicochemical properties and clinical annotation with risk penalties. Risk and Priority are expressed on a 0–100 scale, whereas the Model score is a probability in [0, 1]. The shortlist retains candidates with *T*_max_ *<* 0.3 and Risk *≤* 40, with at most one compound per Bemis–Murcko scaffold.

| Compound | Mechanism class | Clinical annotation | Model score<br>($\bar{p}$) | $T_{\max}$ | Risk | Priority |
| --- | --- | --- | --- | --- | --- | --- |
| Bimatoprost | prostaglandin F2α (FP) | approved (glaucoma) | 0.230 | 0.28 | 0 | 76.5 |
| CE3F4 | EPAC (Epac1)<br>inhibitor | research | 0.540 | 0.25 | 0 | 76.4 |
| Perzinfotel | NMDA-R antagonist | phase 2 (neuropathic<br>pain) | 0.480 | 0.16 | 0 | 75.5 |
| Laurocapram | skin-penetration<br>enhancer | approved (excipient) | 0.263 | 0.24 | 10 | 75.2 |
| NS-309 | SK/IK<sub>Ca</sub> activator | research | 0.411 | 0.18 | 0 | 73.5 |
| 5-Hydroxymethyl-<br>tolterodine | M3 muscarinic<br>antagonist | active metabolite of<br>approved drugs | 0.267 | 0.29 | 18 | 73.0 |

The shortlist includes approved compounds and candidates with clinical development experience, with bimatoprost ranked first. CE3F4 ranked ninth by ensemble probability among the 418 filter-passing records and second by composite priority among candidates retained under the hard exclusion criteria. Its maximum Tanimoto similarity to any molecule in the training screen, active or inactive, is also 0.25. These rankings, together with structural dissimilarity from the training molecules and commercial availability, made this research-stage compound a practical choice for a proof-of-concept assay. Removing exact training-set matches changes neither its ninth position by model score nor the reported shortlist (none of its members is a training-set match). Although the number of filter-passing compounds varies with the predicted-activity cutoff, CE3F4 meets the screening criteria throughout the examined range of [0.15, 0.25]. Score components, exclusion criteria and the cutoff-sensitivity analysis are detailed in Supporting Information, Section S5.

Selecting compounds for antibacterial development requires considering predicted antibacterial activity alongside developability and potential safety liabilities. For example, a cytotoxic anticancer compound may inhibit bacterial growth while also harming mammalian cells. Before triage, the raw ranking includes several cytotoxic antineoplastics, alongside a quinone, a chloromaleimide and a bisbiguanide antiseptic, illustrating why predicted activity is assessed together with these additional criteria.

CE3F4 is a tetrahydroquinoline and a known EPAC1 inhibitor [49–52]. It is listed in the Drug Repurposing Hub as BRD-A47940219, with an ensemble probability of 0.540 and maximum Tanimoto similarity of 0.25 to an active training molecule. It does not trigger the composite ADMET flag, but its predicted probability of drug-induced liver injury is 0.75. This prediction does not activate the flag because the rule requires concomitant hERG risk. The other predicted toxicity probabilities are 0.17 for hERG, 0.02 for ClinTox, 0.28 for Ames mutagenicity and 0.13 for carcinogenicity. During triage, CE3F4 triggered none of the curated structural alerts and received a risk penalty of zero. Because it was the compound taken forward, we also screened it against the complete RDKit implementations of the Brenk and PAINS catalogues. It matches no PAINS patterns and two Brenk patterns. The first is aldehyde, whose SMARTS [C&H1]=O matches the *N*-formyl carbon: the pattern does not exclude an adjacent nitrogen, so it flags this formamide even though a formamide is far less electrophilic than an aldehyde. The second is halogenated_ring_2, which requires three halogens on an aromatic ring; CE3F4 carries two bromines and one fluorine on its ring.

### 3.4 *In vitro* Assay of the CE3F4 Candidate

CE3F4 reduced the growth of *A. baumannii* relative to the vehicle control, with the largest reduction of 57% at 0.25 mM. Higher concentrations produced smaller reductions, yielding a non-monotonic concentration– response profile, as shown in Figure 6.

**Figure 6:**
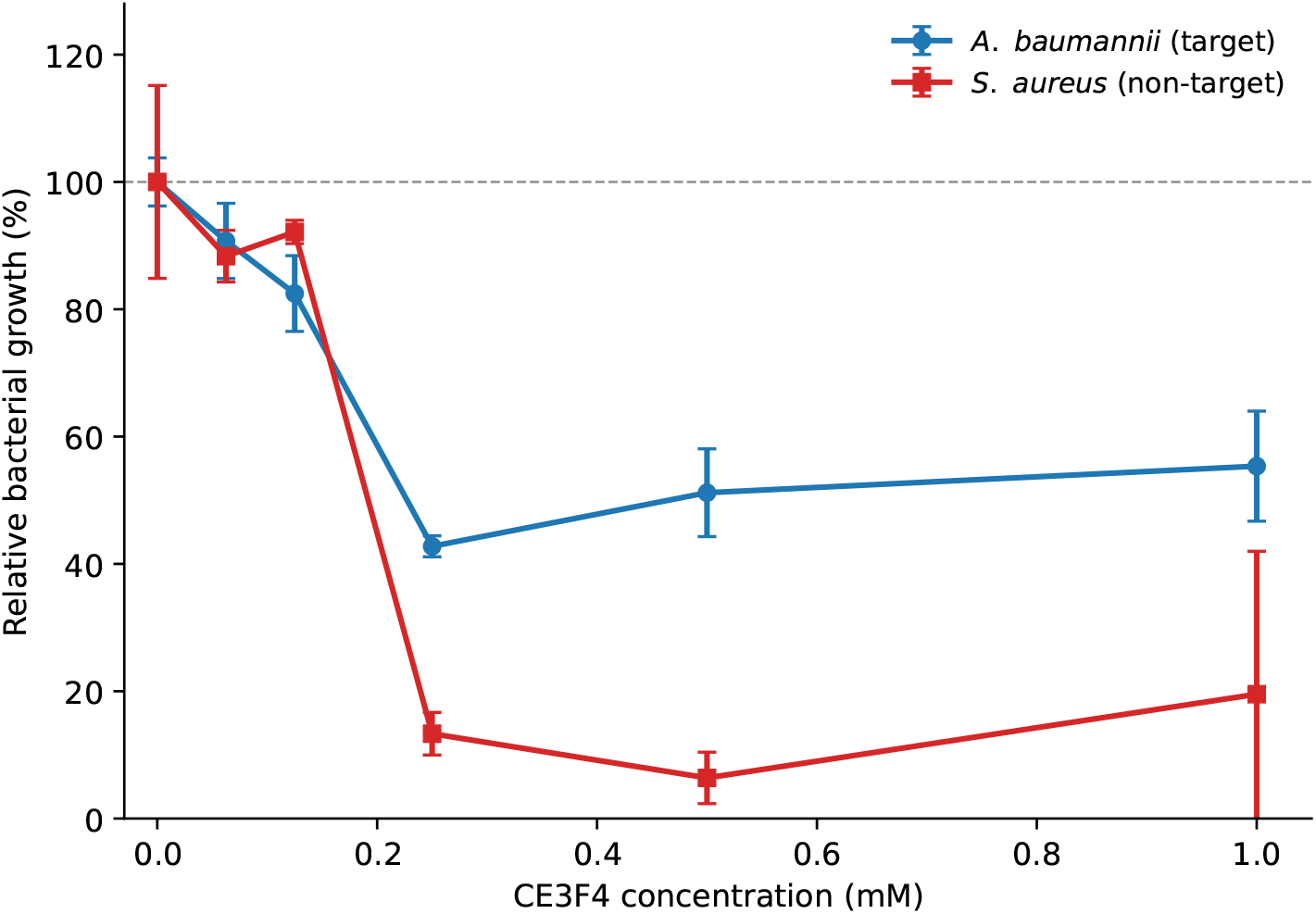
Growth response to CE3F4 in *A. baumannii* NCTC 13305 and the Gram-positive non-target comparator *S. aureus* M/8912. Relative growth was calculated from blank-corrected OD_600_ after 16 h and expressed as a percentage of the matched 5% DMSO vehicle control. Blue circles denote *A. baumannii* and red squares denote *S. aureus*. Points show means and error bars indicate one standard deviation across at least three technical replicate wells from a single preculture per strain. Zero CE3F4 concentration denotes the compound-free vehicle control, whose mean relative growth is marked by the dashed line at 100%. The largest observed growth reductions were 57% at 0.25 mM for *A. baumannii* and 94% at 0.5 mM for *S. aureus*.

Growth reduction was greater in the Gram-positive non-target comparator *S. aureus*, reaching 94% at 0.5 mM, so the effect is not restricted to the Gram-negative target. Only one strain per species was tested and no mammalian-cell cytotoxicity was measured, so no selectivity index is available. These measurements reflect growth at a single 16 h endpoint, so an extended lag phase cannot be fully distinguished from inhibition.

Relative to the 5% DMSO vehicle control, the growth reduction exceeds the well-to-well scatter from 0.125 mM upward for *A. baumannii* and from 0.25 mM upward for *S. aureus*. Because each condition used at least three technical replicate wells from a single preculture, we treat these differences as descriptive and report no inferential test. The reported dispersion therefore reflects assay precision rather than biological variability. At the highest *S. aureus* concentration, the standard deviation exceeds the mean, with relative growth of 19.5 *±* 22.4% at 1 mM, so this point should be read with caution. The vehicle concentration of 5% (v/v) DMSO is higher than usual for whole-cell assays, and higher DMSO concentrations themselves suppressed growth. The concentration of peak effect corresponds to approximately 88 *µ*g/mL for CE3F4’s molecular weight of 351 g/mol. Growth is never fully suppressed within the tested range, so no minimum inhibitory concentration is defined. The inoculum of OD_600_ = 0.05 and the tryptic soy broth medium depart from CLSI broth-microdilution standards, limiting comparability to published MICs. No reference antibiotic was run on the same plates.

CE3F4 was selected through composite prioritization and does not contain the piperazine motif highlighted by the attribution analysis. Whether its lack of a basic amine contributes to limited accumulation in *A. baumannii* remains a hypothesis, informed by accumulation rules derived in *E. coli* [47] and requiring direct uptake measurements.

The non-monotonic response raises the question of how CE3F4 behaves in the assay medium. CE3F4 is poorly water-soluble, and a critical aggregation concentration near 150 *µ*M has been reported for its active *R*-enantiomer, CE3F4R, as discussed in Section 2.7. This reference concentration lies within the range assayed here, below the 0.25 mM concentration of peak growth reduction. In the reference system, CE3F4R forms inert colloidal aggregates that sequester free inhibitor, with dissociation from its EPAC target above approximately 250–350 *µ*M [45]. Such sequestration can limit the availability of dissolved compound and complicate the concentration–response relationship. These observations motivate investigating aggregation in our assay. The assay report notes visible precipitation when CE3F4 was introduced into liquid TSB at 10 mM during preliminary solubility checks, and none at the assayed concentrations, consistent with limited solubility in the medium and with sub-visible aggregation at the higher assayed concentrations. The aggregation threshold for the racemate in TSB with 5% DMSO was not determined.

Testing this possibility with a detergent-based aggregation counter-screen would require controls for effects on both bacteria and compound availability. Non-ionic detergents can alter envelope permeability and antimicrobial susceptibility [53]. Micellar assemblies can also lower the free concentration of hydrophobic compounds, as reported for CE3F4R with Triton X-100 [45]. A loss of signal after detergent addition would therefore need to be interpreted alongside these effects to distinguish aggregation from altered permeability or compound availability.

Chemical stability provides a further possible influence on the response above 0.25 mM. CE3F4 has been reported to undergo hydrolysis at its *N*-formyl group to an inactive product, with a plasma half-life near 40 min [52]. That value was obtained in an amidase-rich matrix, and stability in TSB was not measured. The observed growth reduction therefore motivates follow-up measurements of solubility, aggregation and stability under the assay conditions to determine which processes contribute to the response.

To our knowledge no growth-inhibitory effect on bacteria has previously been reported for CE3F4, a molecule characterized only as an Epac1 inhibitor [49, 50]. This prospective assay connects computational prioritization to an observed bacterial growth response. Further characterization of CE3F4 in the assay medium would help determine the contributions of its solubility, aggregation and stability to the measured response, guiding further experimental work as discussed in Section 4.

The lowest concentration assayed, 62.5 *µ*M, lies above the 50 *µ*M of the primary screen [12] and produced 9% inhibition, within the well-to-well scatter; no measurement was made at 50 *µ*M. The experiment therefore speaks to end-to-end feasibility rather than to transfer of predicted potency.

## 4 Conclusions

This study shows how competitive graph-only prediction can be coupled with evidence about the molecular features supporting predicted antibacterial activity. On the *A. baumannii* screen of Liu et al. [12], this is achieved using atom features and graph connectivity without the approximately 200 additional handcrafted molecular descriptors used by the reference model. Keeping the predictor’s input at the graph level allows attribution analyses to examine changes in atom identity while holding connectivity fixed, without also accounting for a separate molecule-level descriptor input.

Perturbation and restoration tests support the importance of the atoms highlighted by the explanations, both on training actives and external Drug Repurposing Hub molecules. Several recurring motifs show larger perturbation effects in experimentally active than inactive molecules, but their interpretation changes when these effects are compared with equally sized perturbations within the same molecule. Across the examined controls, piperazine provides the most consistent evidence of a recurring motif supporting predicted activity. In parallel, the model’s activity predictions feed a developability and safety assessment that returns a short, chemically diverse candidate list for experimental follow-up. CE3F4, selected from this shortlist, reduced measured bacterial growth under the assay conditions and showed a non-monotonic concentration– response profile. The study thus combines competitive graph-only prediction, evaluation of substructure explanations and candidate prioritization in a documented workflow for prospective antibacterial testing.

The competitive graph-only result motivates evaluation on additional antibacterial screens. Evaluating substructure enrichment with cross-validation models on molecules excluded from their training would test how broadly the chemical rationales transfer. The within-molecule comparison can also be evaluated in other molecular prediction tasks to assess its broader value for chemical interpretation.

For CE3F4, measurements of solubility, aggregation and stability under the assay conditions would guide the choice of concentrations and formulations for further experimental characterization. Prospective testing of further candidates, including MIC, cytotoxicity and subsequent *in vivo* studies where supported by the preceding results, would advance their experimental characterization.

## Data and Software Availability

The training screen is the public dataset of Liu et al. [12], and the prospective library is the Broad Drug Repurposing Hub [6]. Cross-validation fold assignments, candidate tables before and after triage, and summary *in vitro* dose–response measurements are supplied as machine-readable Supporting Information (Data S1–S4). Code and environment specifications are available at https://github.com/AriadneLab/interpretable-gnn-antibacterial. The trained ensemble, its training configuration and the interpretability artifacts underlying the reported figures accompany this submission and reproduce the reported analyses without retraining; they will be deposited in a public archive with a persistent identifier on publication.

## Supporting Information

Model selection and hyperparameter search; cross-validation fold construction and statistical analyses; candidate prioritization definitions and thresholds; supplementary explainability analyses, including held-out faithfulness, occlusion–IG agreement, enrichment robustness, cumulative association and co-occurrence; per-fold results and software versions (PDF); certificate of analysis for CE3F4 (PDF). Cross-validation fold assignments (Data S1), candidates before triage (Data S2), annotated candidates after triage (Data S3), and summary *in vitro* dose–response measurements (Data S4) (CSV).

## Author Information

### Corresponding Author

Cristian Taccioli — Department of Animal Medicine, Production and Health (MAPS), University of Padova, Italy; Department of Computer Science, University of Cambridge, UK;

### Authors

Salvatore Romano — Department of Electrical, Electronic and Computer Engineering, University of Catania, Italy; University Campus Bio-Medico of Rome, Italy; Department of Computer Science, University of Cambridge, UK.

Marco Grassia — Department of Electrical, Electronic and Computer Engineering, University of Catania, Italy.

Dario Righelli — Department of Biology and Department of Statistics, University of Padova, Italy; Department of Computer Science, University of Cambridge, UK.

Pietro Liò — Department of Computer Science, University of Cambridge, UK.

Giuseppe Mangioni — Department of Electrical, Electronic and Computer Engineering, University of Catania, Italy.

### Author Contributions

**S.R.** conceptualization, methodology, software, formal analysis, investigation, data curation, visualization, writing — original draft; **M.G.** methodology, software, visualization, writing — review & editing; **D.R.** formal analysis, writing — review & editing; **P.L.** conceptualization, supervision, writing — review & editing; **G.M.** supervision, methodology, writing — review & editing; **C.T.** supervision, validation, resources, writing — review & editing.

### Notes

The authors declare no competing financial interest.

## Acknowledgments

The authors acknowledge the Austrian Institute of Technology (AIT) for performing the *in vitro* validation assays. During the preparation of this work the authors used LLMs for language editing, and code drafting; the authors reviewed and edited all output and take full responsibility for the content of the manuscript.

