## Supporting Information for "Interpreting Graph Neural Networks for Antibacterial Drug Repurposing against *Acinetobacter baumannii*"

#### Contents

**S1 Model Selection and Hyperparameter Search** **S1**

**S2 Cross-Validation Fold Construction** **S3**

**S3 Statistical Analysis of the Benchmark Comparison** **S3**

**S4 Supplementary Explainability Analyses** **S4**

**S5 Candidate Prioritization and Shortlist** **S8**

**S6 *In vitro* Dose–Response Data** **S12**

**S7 Certificate of Analysis for CE3F4** **S13**

**S8 Machine–Readable Data Files** **S15**

#### S1 Model Selection and Hyperparameter Search

We selected the architecture and training hyperparameters with Optuna [1], running 208 completed trials (of 250 sampled; 42 pruned) across the four GNN backbones (GIN, GINE, GAT, GATv2) and the hyperparameters listed in Table S1. The search optimizes validation ROC-AUC on a single stratified 80/10/10 split. Because performance on an individual split has high-variance, we re-evaluated the five highest-ranking configurations with the highest validation ROC-AUC using the full 10-fold $\times$ 10-model ensemble protocol described in the main text (Table S2). The ranking changes substantially: the best configuration on the single split (trial 188, GINE) falls to last under cross-validation ($0.799 \pm 0.027$, a drop of 0.046), which is exactly why single-split selection is unreliable. Trial 210 (GIN, $K$=4, $d$=256, sum pooling) is the best under cross-validation on both ROC-AUC ($0.819 \pm 0.031$) and PR-AUC ($0.370 \pm 0.060$).

The selected configuration uses a OneCycle learning-rate schedule, which first increases the rate to a maximum of $3.98 \times 10^{-4}$ and then progressively decreases it. The selected Optuna trial had a search budget of 171 epochs, increased to a maximum of 256 epochs for cross-validation and final ensemble training.

Table S1: Optuna search space. Inter-layer batch normalization was fixed off (the two-layer MLP inside each GIN/GINE layer always uses batch normalization); attention-specific hyperparameters (heads, head concatenation) apply to GAT/GATv2 only.

| **Hyperparameters** | **Search space** |
|---|---|
| Backbone | GIN, GINE, GAT, GATv2 |
| Message-passing layers $K$ | $[3, 7]$ |
| Hidden dimension $d$ | $\{64, 128, 256, 512\}$ |
| Pooling | mean, max, sum, concatenated mean and max |
| MLP-head layers | $[1, 3]$ |
| MLP-head hidden dim | $\{64, 128, 256\}$ |
| Dropout | $[0.0, 0.5]$ |
| Residual connections | yes / no |
| Batch normalization | off (fixed) |
| Optimizer | Adam, AdamW |
| Learning rate | log-uniform $[10^{-4}, 10^{-2}]$ |
| Weight decay | log-uniform $[10^{-6}, 10^{-3}]$ |
| LR schedule | none, cosine, plateau, OneCycle |
| Batch size | $\{32, 64, 128, 256\}$ |
| Epochs | $[50, 200]$ |
| Class-imbalance strategy | none, class-weighted, focal |
| Focal $\gamma$ (if focal) | $[0.5, 3.0]$ |
| Attention heads (GAT/GATv2) | $\{2, 4, 8\}$ |
| Concatenate heads (GAT/GATv2) | yes / no |

Table S2: Five configurations with the highest validation ROC-AUC during the Optuna search, re-evaluated using 10-fold cross-validation with a 10-model ensemble in each fold. Rows remain ordered by their validation ROC-AUC on the fixed 80/10/10 partition used during optimization. Cross-validation values are mean ± SD across the ten held-out folds. **Bold** is the configuration selected for deployment.

| **Single-split rank** | **Trial** | **Arch** | **Layers / Hidden** | **Val ROC**$^{\dagger}$ | **CV ROC-AUC** | **CV PR-AUC** |
|---|---|---|---|---|---|---|
| 1 | 188 | GINE | 4 / 128 | 0.845 | $0.799 \pm 0.027$ | $0.318 \pm 0.062$ |
| 2 | 186 | GIN | 5 / 128 | 0.841 | $0.818 \pm 0.030$ | $0.354 \pm 0.068$ |
| 3 | 232 | GIN | 3 / 256 | 0.840 | $0.807 \pm 0.031$ | $0.354 \pm 0.047$ |
| **4** | **210** | **GIN** | **4 / 256** | **0.839** | $\mathbf{0.819 \pm 0.031}$ | $\mathbf{0.370 \pm 0.060}$ |
| 5 | 240 | GIN | 3 / 512 | 0.839 | $0.808 \pm 0.028$ | $0.351 \pm 0.057$ |

$^{\dagger}$Single stratified 80/10/10 split used during search.

**Software** Models and interpretability were implemented in Python 3.10 with PyTorch 2.4.1, PyTorch Geometric 2.5.3, Captum 0.9.0, RDKit 2023.09, scikit-learn 1.7, NumPy 1.26 and SciPy 1.15; the hyperparameter search used Optuna 4.9. The Chemprop baseline and the ADMET-AI predictions were run in their own environments. Exact versions and the full conda environment are released with the code.

### S2 Cross-Validation Fold Construction

For both split protocols, the dataset is partitioned into 10 bins $B_1, \ldots, B_{10}$. For fold $i$, bin $B_i$ is the test set, bin $B_{(i \bmod 10)+1}$ is the validation set (used only for early stopping and checkpoint selection), and the remaining eight bins form the training set, giving an 80/10/10 split per fold. In the random protocol the 7 684 molecules are shuffled once and assigned round-robin to the bins; in the scaffold protocol, Bemis–Murcko scaffolds are greedily packed into the bins (largest first, with each scaffold assigned to the currently smallest bin) so that no scaffold spans more than one bin. Within each fold, $M = 10$ models with distinct seeds are trained and their predicted probabilities are averaged. Because both models are scored on the same fold assignments, the per-fold values (Table S3) are directly paired; this is the basis for the paired significance tests reported below in Section S3. Low-PR folds (e.g., fold 7 under the random split) contain few or hard-to-rank actives, which motivates the multi-fold protocol and the reporting of PR-AUC alongside ROC-AUC.

Table S3: Per-fold test ROC-AUC and PR-AUC for the selected GIN configuration (trial 210) and the Chemprop baseline, on identical folds under both cross-validation protocols (10-fold, 10-model ensemble).

| Split | Model | Metric | 1 | 2 | 3 | 4 | 5 | 6 | 7 | 8 | 9 | 10 | Mean±SD |
|---|---|---|---|---|---|---|---|---|---|---|---|---|---|
| Random | GIN | ROC-AUC | 0.813 | 0.843 | 0.834 | 0.831 | 0.849 | 0.825 | 0.755 | 0.844 | 0.765 | 0.830 | $0.819 \pm 0.031$ |
| Random | Chemprop | ROC-AUC | 0.798 | 0.803 | 0.788 | 0.831 | 0.834 | 0.816 | 0.702 | 0.835 | 0.764 | 0.802 | $0.797 \pm 0.038$ |
| Random | GIN | PR-AUC | 0.405 | 0.391 | 0.378 | 0.386 | 0.445 | 0.448 | 0.240 | 0.325 | 0.307 | 0.378 | $0.370 \pm 0.060$ |
| Random | Chemprop | PR-AUC | 0.380 | 0.424 | 0.383 | 0.413 | 0.453 | 0.375 | 0.244 | 0.334 | 0.300 | 0.438 | $0.374 \pm 0.062$ |
| Scaffold | GIN | ROC-AUC | 0.700 | 0.847 | 0.815 | 0.740 | 0.798 | 0.829 | 0.803 | 0.729 | 0.812 | 0.756 | $0.783 \pm 0.046$ |
| Scaffold | Chemprop | ROC-AUC | 0.695 | 0.810 | 0.772 | 0.722 | 0.785 | 0.805 | 0.707 | 0.706 | 0.804 | 0.806 | $0.761 \pm 0.046$ |
| Scaffold | GIN | PR-AUC | 0.249 | 0.340 | 0.433 | 0.151 | 0.344 | 0.424 | 0.279 | 0.249 | 0.313 | 0.332 | $0.311 \pm 0.080$ |
| Scaffold | Chemprop | PR-AUC | 0.255 | 0.442 | 0.250 | 0.169 | 0.299 | 0.398 | 0.175 | 0.281 | 0.363 | 0.400 | $0.303 \pm 0.090$ |

### S3 Statistical Analysis of the Benchmark Comparison

We assess the GIN-versus-Chemprop comparison using the ten paired fold differences for each metric and split protocol (Table S3). Paired $t$-tests and Wilcoxon signed-rank tests assess differences, and 95% confidence intervals based on the Student's $t$ distribution summarize uncertainty in the mean paired difference. These intervals use the sample standard deviation of the paired differences, whereas the descriptive standard deviations of model performance across folds use the population convention. Holm correction is applied across the four paired $t$-tests. Under random splitting, the ROC-AUC difference is +0.022, with higher GIN performance in 9/10 folds and unadjusted $p < 0.01$ for both tests. Under scaffold splitting, the difference is also +0.022, with higher GIN performance in 9/10 folds, but neither test reaches $\alpha = 0.05$. Holm-adjusted $p$-values are 0.025 and 0.27 for random- and scaffold-split ROC-AUC, respectively. Neither split shows a statistically detectable PR-AUC difference.

Since non-significance alone does not establish equivalence, we also apply two one-sided tests (TOST) [2] to the paired differences with a specified margin of ±0.05 AUC units. The analysis supports equivalence of the mean paired difference within this margin for random-split ROC-AUC ($p = 0.001$), random-split PR-AUC ($p = 0.001$) and scaffold-split ROC-AUC ($p = 0.018$). Scaffold-split PR-AUC remains inconclusive ($p = 0.077$), with a 90% confidence interval of $[-0.042, +0.058]$ extending beyond the margin. However, the folds share training data, so all of these tests and intervals, including TOST, use an independence approximation and may underestimate uncertainty [3]. Their interpretation is also subject to the model-selection and unequal-tuning limitations discussed in the main text.

For context, removing molecular descriptors from the D-MPNN in the reference study reduced its ROC-AUC from 0.792 to 0.756 [4]. Our comparison evaluates a different graph-only modeling workflow against the descriptor-augmented baseline, so it cannot attribute the observed performance differences solely to architecture or descriptor use.

### S4 Supplementary Explainability Analyses

**Full substructure-enrichment results** Table S4 reports every substructure tested for enrichment among the actives. Raw importance is the decrease in the model's predicted activity probability when all atoms of the motif are carbonized together. The table reports its mean among active carriers, while significance is assessed by comparison with inactive carriers containing the same motif. The baseline-controlled excess $\Delta P_{\text{exc}}$ compares the motif's perturbation effect with the mean effect of $R = 3$ equally sized random atom sets within the same molecule, holding the molecule and number of perturbed atoms fixed. Significance is a one-sided Mann–Whitney $U$ test with Benjamini–Hochberg FDR correction. Several motifs are strongly enriched in raw importance, including the bicyclic thioether (cephem-like) core and pyridine, but only piperazine survives the within-molecule baseline control ($\Delta P_{\text{exc}} = 0.25$, $q = 7 \times 10^{-8}$); the remaining motifs, including the benzene negative control, do not ($q \geq 0.05$). Piperazine is thus the only significantly enriched motif under the primary baseline-controlled analysis, and the robustness checks below assess how consistently this model-reliance signal is supported.

Table S4: Substructure enrichment among active relative to inactive motif carriers, ranked by mean raw importance among active carriers. $n$ is the number of active carriers, raw importance is their mean occlusion $\Delta P$, and $\Delta P_{\text{exc}}$ is their mean excess importance relative to equally sized random atom sets within each molecule. $q_{\text{raw}}$ and $q_{\text{ctrl}}$ are Benjamini–Hochberg-adjusted $p$-values from one-sided Mann–Whitney $U$ tests comparing active and inactive carriers.

| **Substructure** | **SMARTS** | $n$ | **Raw** $\Delta P$ | $q_{\text{raw}}$ | $\Delta P_{\text{exc}}$ | $q_{\text{ctrl}}$ |
|---|---|---|---|---|---|---|
| Bicyclic thioether (cephem-like) | `N[C@@H]1CN2C=CCS[C@H]12` | 12 | 0.491 | $4 \times 10^{-4}$ | 0.261 | 0.058 |
| Piperazine | `C1CNCCN1` | 40 | 0.367 | $2 \times 10^{-10}$ | **0.251** | $\mathbf{7 \times 10^{-8}}$ |
| 4-Aminotetrahydropyran | `NC1CCOCC1` | 19 | 0.208 | $6 \times 10^{-3}$ | 0.132 | 0.358 |
| 2-Aminothiazole | `Nc1nccs1` | 9 | 0.180 | $6 \times 10^{-3}$ | 0.004 | 0.803 |
| Pyridine | `c1ccncc1` | 60 | 0.134 | $9 \times 10^{-11}$ | −0.004 | 0.358 |
| Benzene (control) | `c1ccccc1` | 60 | 0.000 | 1.000 | −0.092 | 1.000 |

**Robustness of the substructure enrichment** We stress-tested the within-molecule baseline control along the axes most likely to affect the resulting enrichment estimates (Table S5); every analysis uses the deployed ensemble's attributions and requires no retraining. (i) Number of random fragments. Piperazine's excess importance remains stable as the random baseline is enlarged from $R = 3$ to $R = 100$: the Monte-Carlo error of the baseline falls from 0.085 to 0.014 while the excess ($\approx 0.24$) and its significance ($q \approx 2 \times 10^{-8}$) are similar across these runs; the $R = 3$ estimate reproduces Table S4 to within this Monte-Carlo error. (ii) Connected null. Restricting the null to equally sized connected subgraphs leaves piperazine strongly enriched ($q = 4 \times 10^{-7}$). (iii) Scale-free normalization. Under a within-molecule rank statistic that discards magnitude entirely, the effect remains directional (piperazine outranks 79% of its random fragments in actives versus 58% in inactives) but is not individually significant after FDR correction under this magnitude-free control. (iv) Motif discovery/test split. Splitting the 480 actives into a discovery half (used only to nominate substructures) and a disjoint held-out half (used only to test), piperazine remains strongly enriched on molecules that played no part in its selection ($\Delta P_{\text{exc}} = 0.26$, 21 carriers, $q = 9 \times 10^{-7}$). This separates motif nomination from enrichment testing, while both groups remain part of the deployed ensemble's training data. (v) Scaffold diversity. The 40 active piperazine carriers span 27 distinct Bemis–Murcko scaffolds, with the most frequent scaffold accounting for 12% of carriers.

The bicyclic thioether core is borderline in the primary $R = 3$ analysis ($q = 0.058$) but significant in the sensitivity rerun with $R = 100$ ($q = 0.005$). Its estimate rests on only 12 active carriers and is not significant under the rank statistic. We therefore emphasize piperazine as the most consistently supported motif under the controls varying the number and connectivity of random atom sets, while treating the thioether as a weaker, secondary signal.

**Held-out faithfulness** Figure S1 repeats the deletion/insertion analysis of the main-text faithfulness figure on the top-scoring Drug Repurposing Hub molecules (absent from the training set; exact canonical-

Table S5: Sensitivity of the baseline-controlled piperazine enrichment to the within-molecule random null. $\Delta P_{\mathrm{exc}}$ is the mean excess importance over $R$ random equal-size fragments per carrier, reported separately for active and inactive piperazine carriers; MC error is the Monte-Carlo standard error of that baseline; $q_{\mathrm{ctrl}}$ is the Benjamini–Hochberg-adjusted $p$-value from the one-sided Mann–Whitney $U$ comparison of active and inactive carriers, as in Table S4.

| | | $\Delta P_{\mathrm{exc}}$ | | | $q_{\mathrm{ctrl}}$ | |
|---|---|---|---|---|---|---|
| **Random null** | $R$ | **active** | **inactive** | **MC error** | **piperazine** | **thioether** |
| Disconnected atoms | 3 | 0.232 | 0.024 | 0.085 | $1 \times 10^{-8}$ | 0.050 |
| Disconnected atoms | 50 | 0.242 | 0.024 | 0.019 | $2 \times 10^{-8}$ | 0.013 |
| Disconnected atoms | 100 | 0.238 | 0.023 | 0.014 | $1 \times 10^{-8}$ | 0.005 |
| Connected subgraph | 50 | 0.197 | 0.027 | 0.020 | $4 \times 10^{-7}$ | 0.036 |

SMILES matches removed). The occlusion/IG-versus-random separation is at least as large as on the training actives. These results extend support for the attribution rankings under the evaluated perturbations to molecules absent from training.

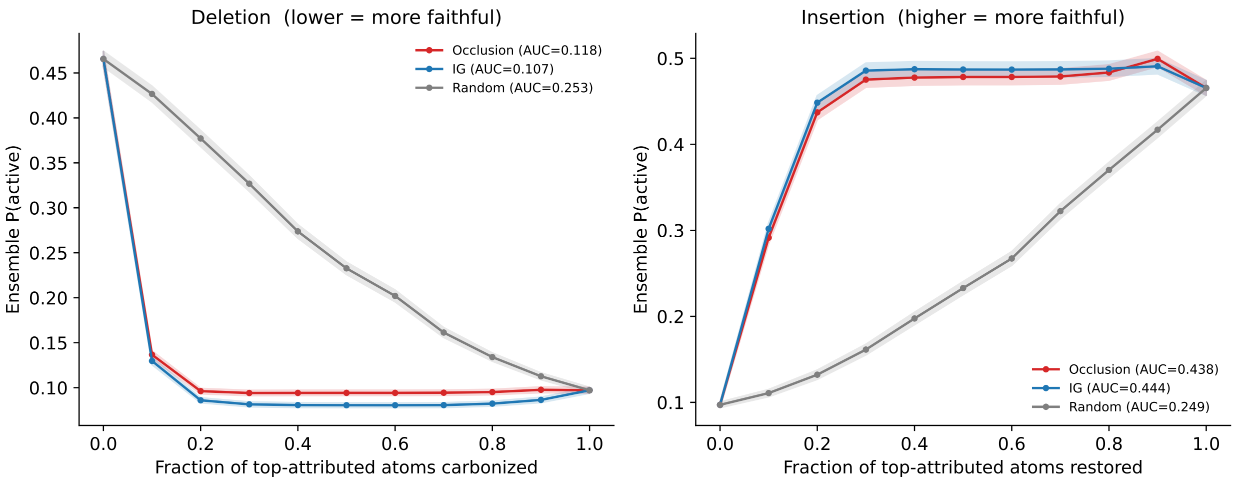


Figure S1: Held-out faithfulness on top-scoring Drug Repurposing Hub molecules absent from training set. Deletion (left) progressively replaces atom identities with carbon in decreasing attribution order, whereas insertion (right) progressively restores the original atom identities from the carbon-reference graph.

**Occlusion–IG agreement** Figure S2 shows the per-molecule Spearman correlation between the atom-level rankings produced by the two attribution methods (median $\rho = 0.72$), providing a measure of inter-method agreement.

**Cumulative association and co-occurrence** We additionally examined whether the number of enriched substructures carried by a molecule is associated with experimental activity or model score. Molecules carrying more of these substructures tend to have higher ensemble scores in the Drug Repurposing Hub (Spearman $\rho = 0.09$, $p = 1.4 \times 10^{-13}$) and a higher frequency of experimental activity in the training set ($\rho = 0.06$, $p = 4.5 \times 10^{-7}$; Figure S3). Both associations are weak, and the groups carrying multiple motifs are small, so these results describe cumulative association rather than demonstrating additive pharmacophoric contributions.

The substructures also co-occur in active molecules more often than expected under independence. The strongest off-diagonal associations are between the bicyclic thioether core and 2-aminothiazole (lift 26.7, $n = 6$), consistent with their joint occurrence in cephalosporin-like structures, and between piperazine and

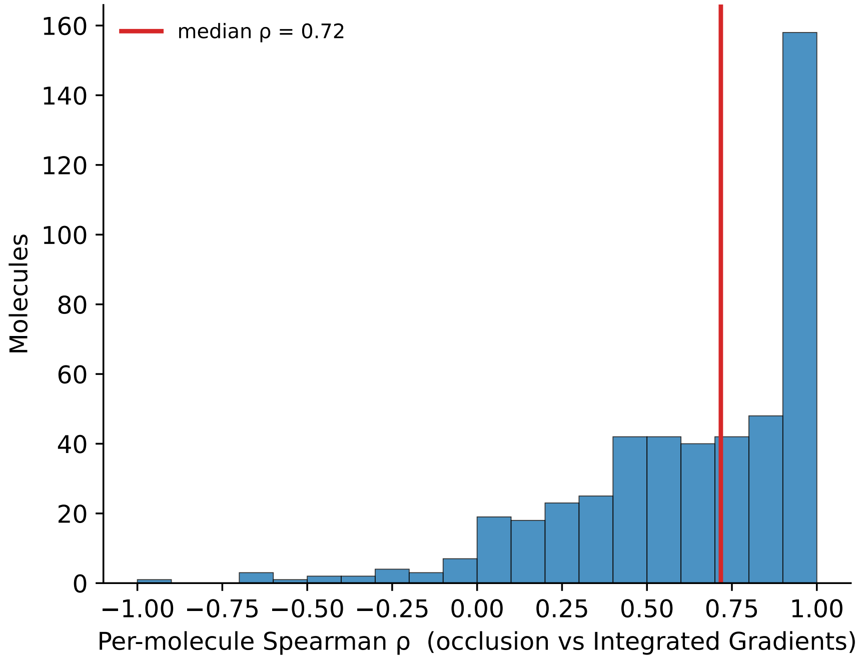


Figure S2: Distribution of per-molecule Spearman correlations between occlusion and Integrated Gradients atom rankings across training actives. The red vertical line marks the median correlation, $\rho = 0.72$. Positive values indicate agreement in the relative ranking of atoms, whereas negative values indicate opposing rankings.

pyridine (lift 2.8, $n = 18$), consistent with their joint occurrence in fluoroquinolone-like structures (Figure S4). The 418 filter-passing repurposing records also contain at least one of these motifs somewhat more frequently than the full Drug Repurposing Hub (28.7% versus 24.0%; Figure S5). These co-occurrence and prevalence comparisons provide descriptive context for motifs that, with the exception of piperazine, do not remain significant under the primary within-molecule control (Table S4).

**Population-level attributions** To extend the characterization of model reliance beyond the training-actives analysis, we examined motif effects across all 6 642 curated Drug Repurposing Hub compounds. For each compound, we jointly carbon-substituted the atoms matching each motif and recorded the resulting change in ensemble probability, $\Delta P$, using the operation defined in Section 2.6. This population includes training-set overlaps and complements the separate faithfulness evaluation on molecules absent from training. Among the 407 piperazine-bearing compounds, the Spearman correlation between the intact score and $\Delta P_{\text{piperazine}}$ is +0.84. For Figure S6, compounds are grouped into probability intervals of width 0.1. Piperazine prevalence increases overall from 3.9% in the lowest interval to 56% in the highest, with fluctuations between intermediate intervals. The effect extends to the broader class of basic amines that piperazine exemplifies (Spearman +0.47 over all compounds, +0.64 among the 3 677 carriers). Protonatable amines have been associated with accumulation in Gram-negative bacteria [5]; we note the correspondence as chemical context only, since these correlations relate model predictions to perturbation effects and no uptake was measured here.

Halogen perturbations show a weaker association (Spearman +0.28), with mean $\Delta P$ decreasing to approximately 0.01 in the highest probability interval. A partial Spearman correlation controlling for the halogen perturbation effect leaves the basic-amine association at +0.45.

The within-motif correlations summarize how perturbation effects vary with the intact model score under a fixed motif definition. These analyses characterize the deployed model's behavior across the curated library, rather than establishing experimental antibacterial activity. Table S6 illustrates the attribution patterns on selected named compounds. The higher-scoring reference compounds show their largest tested motif-level

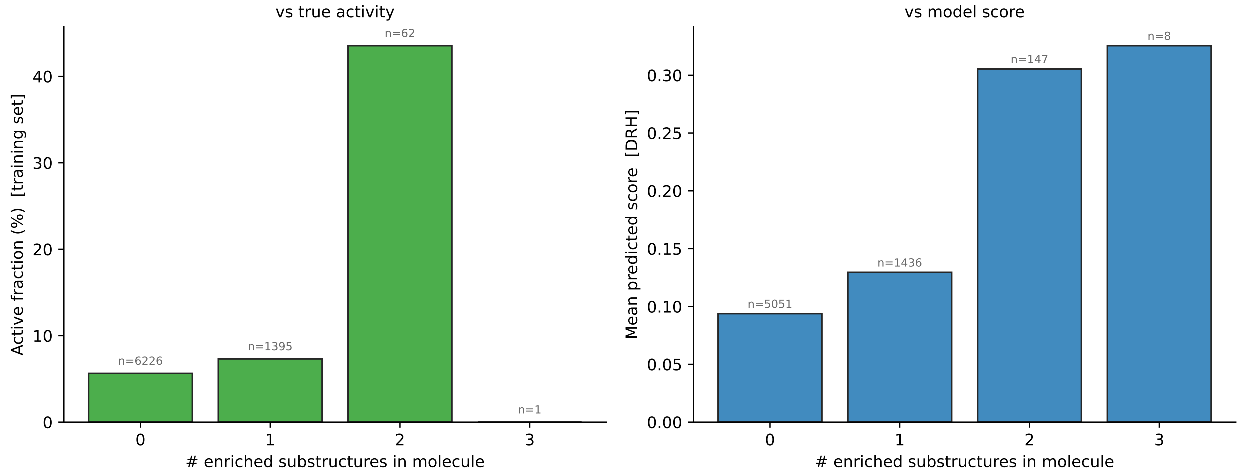


Figure S3: Experimental activity and prediction score by motif count. Left, fraction of active training molecules as a function of the number of enriched motifs carried by each molecule. Right, mean ensemble score across Drug Repurposing Hub compounds grouped by the same motif count. Labels above the bars report the number of molecules in each group.

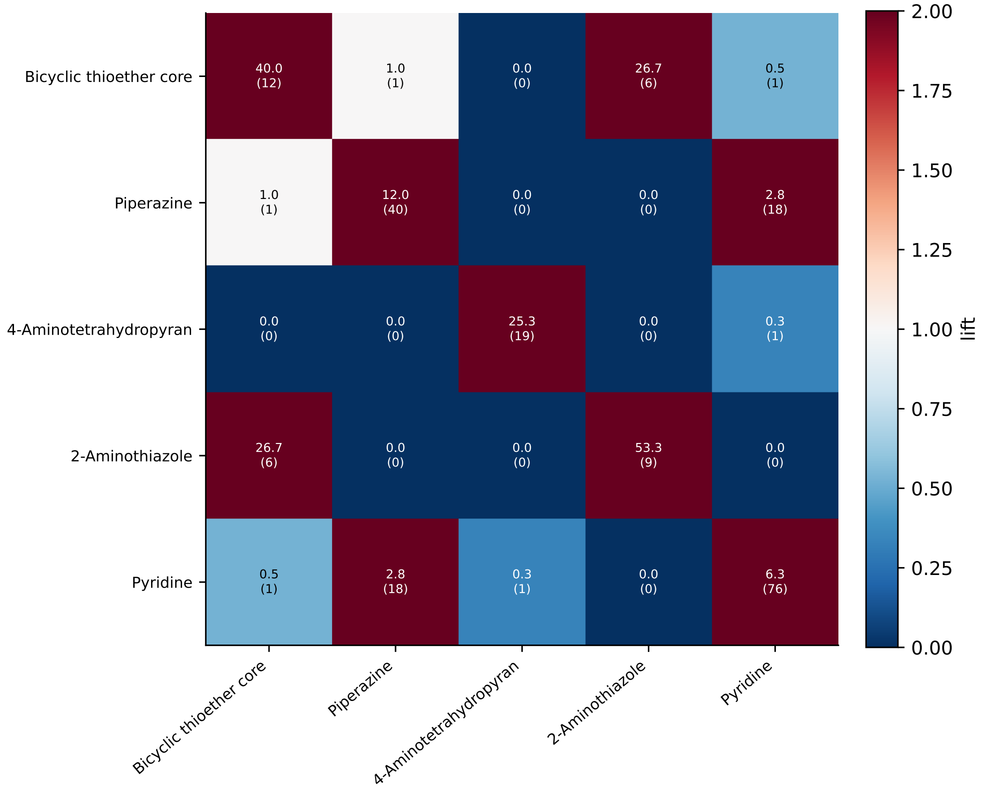


Figure S4: Pairwise co-occurrence lift among candidate motifs in active molecules. Lift is the observed joint frequency divided by the expected joint frequency under independence, and joint counts are shown in parentheses.

perturbation effects for basic amines, including piperazine in dovitinib, whereas CE3F4 shows its largest

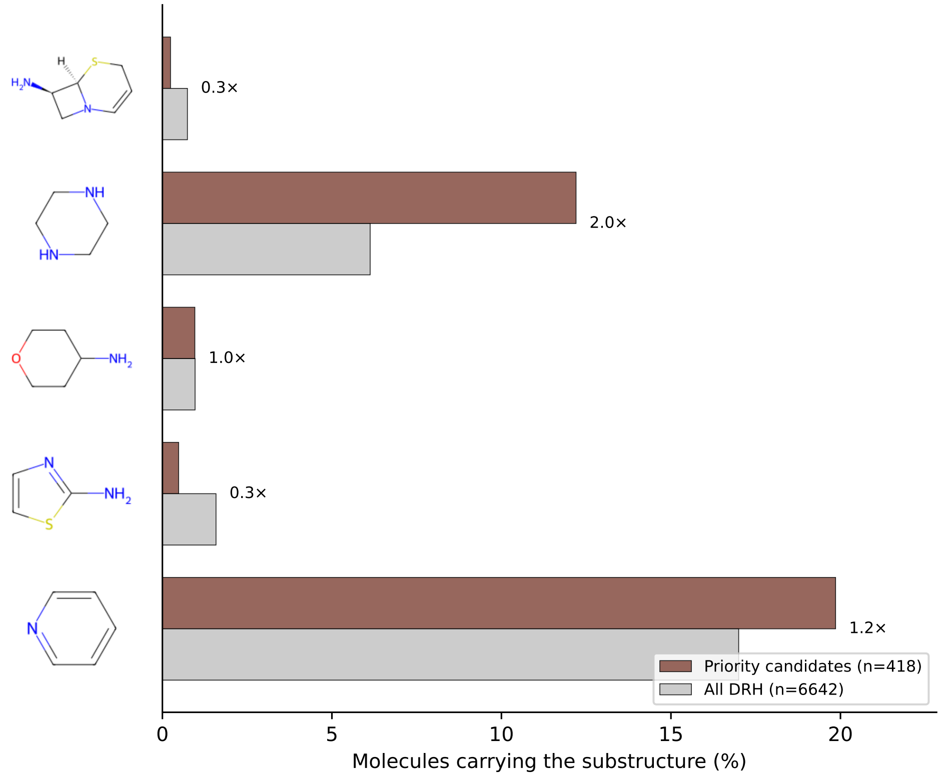


Figure S5: Frequency of each candidate motif among the 418 Drug Repurposing Hub records passing the predicted-activity and similarity filters and among all 6 642 curated Drug Repurposing Hub compounds. Labels report the ratio of the motif frequency in the filter-passing set to that in the full library.

tested effect for halogen atoms. These examples are illustrative and do not establish that the reported motifs determine experimental antibacterial activity.

Table S6: Per-candidate attribution summaries for representative higher- and lower-scoring Drug Repurposing Hub compounds. $\bar{p}$ is the ensemble score. The reported dominant substructure is the tested motif class with the largest positive carbon-reference perturbation effect for that compound; $\Delta P$ is the corresponding decrease in $\bar{p}$. The designation does not imply that the motif is a biologically validated activity determinant.

| **Compound** | **Role** | $\bar{p}$ | **Dominant substructure** ($\Delta P$) |
|---|---|---|---|
| eravacycline | top-ranked (known antibiotic) | 0.858 | basic amine (+0.47) |
| NSC-663284 | ranking control | 0.626 | basic amine (+0.35) |
| IT1t | ranking control | 0.577 | basic amine (+0.17) |
| dovitinib | risk control | 0.573 | **piperazine** (+0.46) |
| irinotecan | risk control | 0.566 | basic amine (+0.41) |
| CE3F4 | shortlist (*in vitro*) | 0.540 | halogen (+0.39) |

### S5 Candidate Prioritization and Shortlist

Structural alerts were applied using two curated pattern sets comprising 15 reactive-chemistry alerts and 18 nuisance or assay-risk alerts, listed in Table S7. Eight alerts in each set were triggered by at least one screened candidate, while the remaining patterns matched no candidate and are retained in the table for

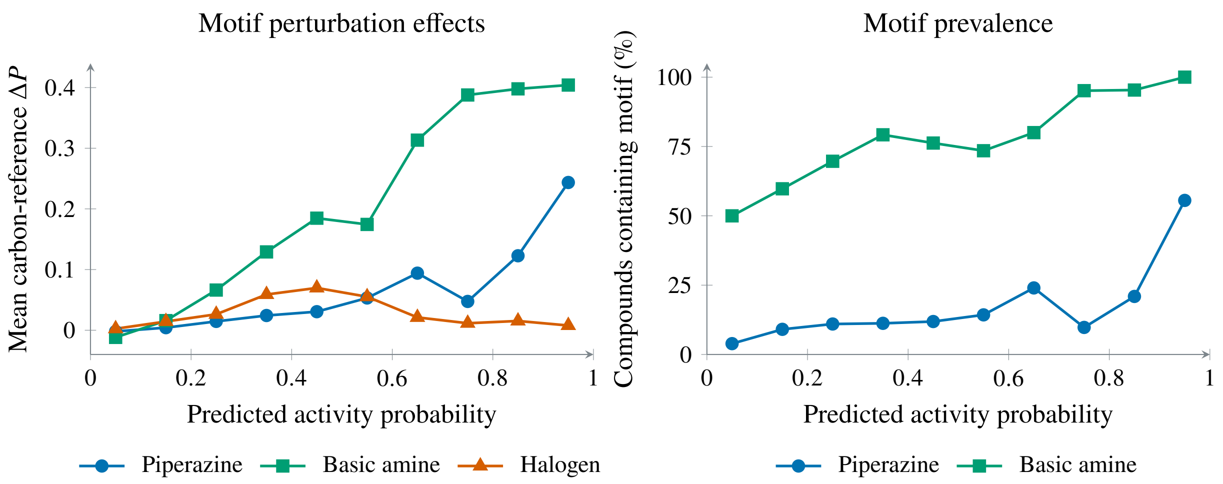


| Interval | 0.0–0.1 | 0.1–0.2 | 0.2–0.3 | 0.3–0.4 | 0.4–0.5 | 0.5–0.6 | 0.6–0.7 | 0.7–0.8 | 0.8–0.9 | 0.9–1.0 |
|---|---|---|---|---|---|---|---|---|---|---|
| Compounds | 4475 | 1257 | 455 | 178 | 101 | 49 | 25 | 41 | 43 | 18 |

Figure S6: Population-level attribution patterns across all 6 642 curated Drug Repurposing Hub compounds, including training-set overlaps. Left, mean carbon-reference $\Delta P$ within probability intervals of width 0.1 after joint carbon substitution of piperazine, basic-amine or halogen atoms. Right, prevalence of each motif within the same intervals. Points are positioned at interval midpoints, and the table below the plots reports the number of compounds in each interval. Piperazine and basic-amine prevalence and perturbation effects generally increase with score, whereas the mean halogen perturbation effect peaks at intermediate scores and declines in the highest intervals.

completeness. These patterns were informed by the literature on reactive and nuisance substructures, but do not correspond to the complete published PAINS [6] or Brenk [7] catalogues. CE3F4 matched none of the 33 curated alerts and received a risk penalty of zero. Its subsequent assessment against the complete PAINS and Brenk catalogues is reported in the main text. Three queries ($\alpha$-halo carbonyl, permanent cation and polycation) matched no candidate as applied, and are listed in the form used to produce the released scores. Rewritten with the halogen as a substituent and with the charged nitrogen quaternary, they would flag one further compound, outside the shortlist, and otherwise duplicate the quaternary-ammonium alert; the shortlist and all reported scores and rankings are unaffected.

**ADMET toxicity flag** Six endpoints are predicted with ADMET-AI. A compound receives a composite toxicity flag if hERG $\geq 0.80$, or ClinTox $\geq 0.70$, or (hERG $\geq 0.50$ and DILI $\geq 0.50$); $LD_{50}$, Ames mutagenicity and carcinogenicity are reported for context. Structural alerts enter the triage separately through the chemical-reactivity and assay-artifact component of the risk penalty, using the curated patterns in Table S7.

**Risk penalty** (0–100) Risk is aggregated over four risk categories via a weighted maximum: (1) cardiotoxicity (QT/torsade signals, regulatory box warnings), (2) systemic/organ toxicity (market withdrawal, hepatotoxicity/cytotoxicity classes and the ADMET flag above), (3) chemical reactivity/assay-artifact risk (reactive and nuisance substructures), and (4) an epistemic no-data penalty for compounds lacking annotation.

**Priority score** Candidates are ranked by a heuristic composite score, normalized to 0–100, with priority $\propto$ $0.55\,a + 0.35\,t - 0.30\,r$. The activity-oriented term $a$ combines the ensemble probability with weight 0.60, continuous structural novelty with weight 0.18, a binary structural-novelty indicator with weight 0.10, and a physicochemical developability proxy with weight 0.12. Continuous novelty is one minus the maximum Tanimoto similarity to an experimentally active training molecule. The binary indicator records whether the candidate passes the structural-novelty criterion. Both novelty components therefore derive from the same

Table S7: Curated structural-alert patterns used in the developability and safety triage. Each alert is encoded as a SMARTS pattern, and each match contributes to the chemical-reactivity or assay-artifact component of the risk penalty. The final column reports the number of screened candidates matching each pattern. Patterns that matched no candidate are retained for completeness.

| **Alert** | **SMARTS** | **Matches** |
|---|---|---|
| *Reactive chemistry* | | |
| Michael acceptor | `[C,c]=[C,c]-[CX3](=O)[O,N,#6]` | 17 |
| Acyl hydrazide | `[CX3](=O)[NX3][NX3]` | 7 |
| Phosphonate ester | `[PX4](=O)([OX2])([OX2])[OX2]` | 6 |
| Epoxide | `C1OC1` | 4 |
| Isothiocyanate | `N=C=S` | 3 |
| Disulfide | `[SX2][SX2]` | 2 |
| Aldehyde | `[CX3H1](=O)[#6]` | 1 |
| Hydroxamic acid | `[CX3](=O)[NH][OH]` | 1 |
| $\alpha$-Halo carbonyl | `[CX4;H1,H2][Cl,Br,I][CX3](=O)` | 0 |
| $\beta$-Lactone | `C1C(=O)OC1` | 0 |
| Acyl halide | `[CX3](=O)[Cl,Br,I,F]` | 0 |
| Aziridine | `C1CN1` | 0 |
| Nitroso | `[NX2]=O` | 0 |
| Sulfonyl halide | `[SX4](=O)(=O)[Cl,Br]` | 0 |
| Vinyl sulfone | `[SX4](=O)(=O)C=C` | 0 |
| *Nuisance / assay risk* | | |
| Long alkyl chain | `CCCCCCCCCC` | 36 |
| Catechol | `c1cc(O)c(O)cc1` | 22 |
| Hydroquinone | `Oc1ccc(O)cc1` | 7 |
| Quaternary ammonium | `[N+](C)(C)(C)` | 7 |
| Hydroxamate metal chelator | `C(=O)NO` | 6 |
| Catechol metal chelator | `[OH]c1ccccc1[OH]` | 4 |
| Nitroaromatic | `c1ccccc1[N+](=O)[O-]` | 3 |
| Quinone | `O=C1C=CC(=O)C=C1` | 1 |
| Alkylidene barbiturate | `C=C1C(=O)NC(=O)NC1=O` | 0 |
| Aryl thiocarbonyl | `c1ccccc1C(=S)` | 0 |
| Azo dye | `c1ccccc1N=Nc2ccccc2` | 0 |
| Ene-rhodanine | `C=C1SC(=S)NC1=O` | 0 |
| Hydroxyphenyl hydrazone | `c1cc(O)ccc1/N=N` | 0 |
| Permanent cation | `[N+;!H0]` | 0 |
| Polycation | `[N+;!H0].[N+;!H0]` | 0 |
| Polycyclic aromatic | `c1ccc2c(c1)cc1ccc3ccccc3c1c2` | 0 |
| Rhodanine | `O=C1CSC(=S)N1` | 0 |
| Sulfonium or phosphonium | `[S+,P+]` | 0 |

similarity measure, and the binary component is constant within the novelty-filtered pool. The translational term $t$ combines clinical readiness and repurposing opportunity, weighted 0.70 and 0.30, respectively. Clinical readiness incorporates safety annotation, evidence of UK/EU approval or use and available human data, whereas repurposing opportunity incorporates non-antibiotic status and the intellectual-property opportunity annotation. Clinical readiness credits human-exposure evidence arising from an approved parent drug or prodrug, so an active metabolite of an approved product is treated as having use evidence. 5-Hydroxymethyl-tolterodine, the active metabolite of both tolterodine and fesoterodine, is credited on this basis; it is not itself marketed as a product.

This term does not include the physicochemical developability proxy used in $a$. The risk penalty $r$ incorporates toxicity and safety information, so safety-related criteria contribute to both $t$ and $r$. These overlapping components are prioritization heuristics rather than independent estimates of candidate quality. The main shortlist retains compounds with risk penalty $\leq 40$ and at most one compound per Bemis–Murcko scaffold. Annotation is retrieved by structure and by name, and candidates for which no record is retrieved receive the no-data penalty rather than clinical credit, so their position is conservative. Fesoterodine is one such case: annotating it as the approved product it is would raise its composite score from 47.0 to 67.4, near rank 13, still outside the shortlist, whose lowest member scores 73.0.

The resulting shortlist (reported in the main text) is dominated by approved or late-stage drugs that rank highly on developability despite modest model scores; by the composite priority score its top entry is bimatoprost, with CE3F4 a close second. CE3F4 carries the highest raw model score within the shortlist but is not the single highest-scoring prediction of the model overall (Table S9); it was advanced to *in vitro* testing as a proof-of-concept, a research-stage, structurally novel and commercially available compound among the top of the shortlist, rather than on its raw prediction alone.

**Sensitivity to the score threshold** The $\bar{p} > 0.2$ activity cut-off is inherited from Liu et al. [4] and sets only the size of the candidate pool, which varies smoothly with it: 750, 418 and 252 compounds pass at $\bar{p} > 0.15$, 0.20 and 0.25 (all with maximum Tanimoto $< 0.3$). Because the ensemble is trained with unweighted cross-entropy, $\bar{p}$ is used as a monotone ranking to size this pool rather than as a calibrated risk estimate, and the shortlist is determined by the composite priority score, not by the raw threshold. The compound taken forward, CE3F4 ($\bar{p} = 0.540$), lies well inside the pool at every threshold in this range, so the experimental decision does not depend on the exact cut-off. At the $\bar{p} > 0.2$ cutoff, Data S2 contains 418 filter-passing records, of which 12 are exact training-set matches and 406 are absent from training. Two of the unmatched records refer to the same compound, cyanocobalamin: they share the InChIKey RMRCNWBMXRMIRW-WZHZPDAFSA-M and differ only by a stereo descriptor on a non-stereogenic phosphorus, which the canonical SMILES retains and the InChI does not. Deduplication is performed on the InChIKey, leaving 405 distinct candidates absent from training. Data S3 contains 417 distinct annotated and scored compounds, including all twelve training matches. Of these, 398 pass the hard exclusion criteria, while 19 are excluded based on curated toxicity classes or known-antimicrobial annotation. The 398 records still include the twelve training-set matches, so this count describes eligibility under the hard exclusion criteria rather than the final prospective population. CE3F4 has the third-highest priority score across all scored records and ranks second after the hard exclusions. None of the twelve exact training matches entered the shortlist.

Three of the screened records are organometallic compounds: bis(maltolato)oxovanadium(IV), mangafodipir, and auranofin. Their metal centres lie outside the atom vocabulary of the featurization and are represented in a single out-of-vocabulary category. The model holds no learned representation for these centres, so the scores assigned to such structures are not interpretable as predictions. They are also the only records whose released scores are not reproduced when recomputed from the released code and checkpoints; the recomputed values fall below the retention threshold, and the difference could not be traced to the featurization, the input structures or the inference procedure. None of the three was shortlisted, and the compound taken forward is unaffected.

**Predicted ADMET of the shortlist** Table S8 gives the six ADMET-AI endpoints for the shortlisted compounds. Classification endpoints (hERG, ClinTox, DILI, Ames, carcinogenicity) are predicted probabilities in $[0, 1]$ (higher = higher risk); $LD_{50}$ (Zhu) is a regression score. CE3F4 does not trigger the composite flag (hERG 0.17, ClinTox 0.02, and hERG $< 0.5$ so the joint hERG/DILI rule does not apply), consistent with its retention in the shortlist; its only elevated endpoint is DILI (0.75). Two other shortlisted compounds do carry an individually elevated endpoint. 5-Hydroxymethyl-tolterodine has hERG 0.84, which fires the composite flag (hERG $\geq 0.80$); it was nonetheless retained because the flag is only one input to the graded 0–100 risk penalty, whose aggregate value stayed low (Risk 18). NS-309 has a high DILI (0.95) but low hERG (0.19), so under our rule, which flags DILI only jointly with hERG $\geq 0.50$, it is not flagged (Risk 0); the rule's lenient treatment of standalone DILI is a limitation of the triage. "Predicted-safe" here therefore denotes low aggregate predicted risk under this rule, not the absence of any elevated endpoint.

**Model ranking versus prioritization** Table S9 lists the fifteen highest-scoring compounds by ensemble probability $\bar{p}$ within the candidate pool passing the predicted-activity and structural-dissimilarity filters, before the developability and safety assessment. This comparison shows how the subsequent assessment reshapes the model ranking within that pool. The highest-scoring compound is eravacycline, a marketed tetracycline-class antibiotic, which is retained as a prediction control and excluded from experimental prioritization as an existing antibiotic. Several other high-scoring compounds are retained only as prediction controls, placed on hold or excluded. CE3F4 ranks ninth by ensemble probability in this pool and combines structural dissimilarity from active training molecules with commercial availability. It does not trigger the composite toxicity flag, although its DILI prediction is elevated and structural alerts are reported in the main text.

Table S9: Top-fifteen prospective candidates by raw ensemble score $\bar{p}$ (before the developability/safety triage), among the structurally novel compounds ($T_{\max} < 0.3$). *Outcome* is the triage disposition: shortlist, hold, control or excluded.

| **Rank** | **Compound** | **Model score ($\bar{p}$)** | $T_{\max}$ | **Outcome** |
|---|---|---|---|---|
| 1 | Eravacycline | 0.858 | 0.25 | Excluded (known antibiotic) |
| 2 | BAX channel blocker | 0.684 | 0.26 | Excluded |
| 3 | NSC-663284 | 0.626 | 0.27 | Control |
| 4 | IT1t | 0.577 | 0.16 | Control |
| 5 | Dovitinib | 0.573 | 0.26 | Control |
| 6 | Chlorisondamine diiodide | 0.568 | 0.21 | Excluded |
| 7 | Irinotecan | 0.566 | 0.19 | Control |
| 8 | RI-1 | 0.556 | 0.29 | Hold |
| 9 | CE3F4 | 0.540 | 0.25 | **Shortlist (tested)** |
| 10 | Semapimod | 0.535 | 0.24 | Hold |
| 11 | TCID | 0.534 | 0.25 | Hold |
| 12 | Alexidine | 0.526 | 0.27 | Hold |
| 13 | Cobimetinib | 0.515 | 0.20 | Hold |
| 14 | Satraplatin | 0.506 | 0.22 | Hold |
| 15 | ALX-40-4C | 0.505 | 0.18 | Hold |

#### S6 *In vitro* Dose–Response Data

Table S10 lists the summary broth-microdilution measurements (mean ± SD over the technical replicate wells) underlying the dose–response curves of the main text: blank-corrected relative growth (% of the 5% DMSO compound-free control) for *A. baumannii* NCTC 13305 (target) and *S. aureus* M/8912 (Gram-positive non-target control), with the corresponding growth inhibition. Each condition is the mean ± SD over $\geq 3$ technical replicate wells dispensed from a single preculture per strain. The tested material was racemic CE3F4 (MedChemExpress HY-108539, lot 279781), for which the supplier's Certificate of Analysis (COA) reports 98.85% purity by HPLC, with $^1$H NMR and MS consistent with the structure. The COA is reproduced in Section S7. Against *A. baumannii* inhibition is modest and non-monotonic, peaking at 0.25 mM (57%) and not increasing further at higher concentrations; against *S. aureus* inhibition is much stronger ($\geq$ 80% from 0.25 mM), confirming that the activity is not selective for the intended Gram-negative target.

#### S7 Certificate of Analysis for CE3F4

Supplier Certificate of Analysis for racemic CE3F4 (MedChemExpress HY-108539, lot 279781), reporting 98.85% purity by HPLC with $^{1}$H NMR and MS consistent with the structure.

MCE®
MedChemExpress

### Certificate of Analysis



## CE3F4

**Cat. No.:** HY-108539

**CAS No.:** 143703-25-7

**Batch No.:** 279781

**Chemical Name:** 1(2H)-Quinolinecarboxaldehyde, 5,7-dibromo-6-fluoro-3,4-dihydro-2-methyl-

##### PHYSICAL AND CHEMICAL PROPERTIES

**Molecular Formula:** $C_{11}H_{10}Br_2FNO$

**Molecular Weight:** **351.01**

**Storage:** -20°C, stored under nitrogen
* In solvent : -80°C, 6 months; -20°C, 1 month (stored under nitrogen)

**Chemical Structure:**

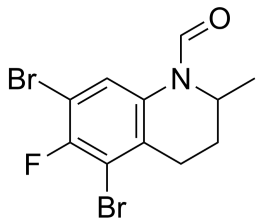


##### ANALYTICAL DATA

**Appearance:** White to off-white (Solid)

**$^1$H NMR Spectrum:** Consistent with structure

**MS:** Consistent with structure

**Purity (HPLC):** **98.85%**

**Conclusion:** The product has been tested and complies with the given specifications.

**Caution: Product has not been fully validated for medical applications. For research use only.**

Address: 1 Deer Park Dr, Suite F, Monmouth Junction, NJ 08852, USA

Table S8: ADMET-AI predictions for the shortlisted compounds. Values are predicted probabilities in $[0,1]$ except $LD_{50}$, which is the Zhu regression score. Bold values meet the component thresholds hERG $\geq 0.80$, ClinTox $\geq 0.70$ or DILI $\geq 0.50$. An elevated DILI prediction triggers the composite toxicity flag only when hERG is also $\geq 0.50$, whereas hERG $\geq 0.80$ or ClinTox $\geq 0.70$ triggers the flag individually.

| **Compound** | **hERG** | **ClinTox** | **DILI** | **$LD_{50}$ (Zhu)** | **Ames** | **Carcinogen.** |
|---|---|---|---|---|---|---|
| CE3F4 | 0.17 | 0.02 | **0.75** | 2.43 | 0.28 | 0.13 |
| Bimatoprost | 0.75 | 0.21 | 0.10 | 2.52 | 0.25 | 0.05 |
| Perzinfotel | 0.06 | 0.32 | **0.62** | 1.55 | 0.43 | 0.28 |
| Laurocapram | 0.58 | 0.02 | 0.12 | 1.75 | 0.03 | 0.04 |
| NS-309 | 0.19 | 0.09 | **0.95** | 2.70 | 0.64 | 0.34 |
| 5-Hydroxymethyl-tolterodine | **0.84** | 0.22 | 0.08 | 2.68 | 0.17 | 0.17 |

Table S10: *In vitro* dose–response (summary). Relative growth is mean ± SD (% of the 5% DMSO control) over $\geq 3$ technical replicate wells; inhibition $= 100 -$ growth.

| **CE3F4 (mM)** | *A. baumannii* (target) | | *S. aureus* (non-target) | |
|---|---|---|---|---|
| | **Growth (%)** | **Inhib. (%)** | **Growth (%)** | **Inhib. (%)** |
| 0 (5% DMSO) | $100.0 \pm 3.8$ | 0 | $100.0 \pm 15.2$ | 0 |
| 0.0625 | $90.7 \pm 5.9$ | 9 | $88.3 \pm 4.1$ | 12 |
| 0.125 | $82.5 \pm 5.9$ | 18 | $92.1 \pm 1.8$ | 8 |
| 0.25 | $42.8 \pm 1.7$ | 57 | $13.3 \pm 3.3$ | 87 |
| 0.5 | $51.2 \pm 6.9$ | 49 | $6.4 \pm 4.0$ | 94 |
| 1.0 | $55.4 \pm 8.6$ | 45 | $19.5 \pm 22.4$ | 80 |

### S8 Machine–Readable Data Files

The following files accompany this Supporting Information and the public repository. The activity labels of the training screen are not redistributed here: they are the published dataset of Liu et al. [4] and should be obtained from that source. The screen structures appear in Data S1 because the fold assignments cannot be reproduced without them.

- **Data S1** — cross-validation fold assignments: canonical SMILES mapped to their random-split and Bemis–Murcko scaffold-split fold indices (1–10), reproducing the exact partitions used throughout (activity labels omitted; taken from Liu et al. [4]) (CSV).
- **Data S2** — model priority candidates before triage: the 418 Drug Repurposing Hub records passing the filters (ensemble score $> 0.2$, maximum Tanimoto $< 0.3$) with Broad id, SMILES, ensemble score, structural-novelty flag and a flag marking the twelve exact training-set matches; the 406 unflagged rows represent 405 distinct compounds absent from training, and the file provides the raw model ranking of Table S9 (CSV).
- **Data S3** — 417 annotated and scored candidate records, with the six ADMET-AI endpoints, curated structural-alert matches (Table S7), risk penalty, composite priority score and selection outcome (CSV).
- **Data S4** — summary *in vitro* dose–response measurements underlying Table S10 (CSV).
